# Distinct cDC subsets co-operate in CD40 agonist response while suppressive microenvironments and lack of antigens subvert efficacy

**DOI:** 10.1101/2021.12.25.474021

**Authors:** Aleksandar Murgaski, Máté Kiss, Helena Van Damme, Daliya Kancheva, Isaure Vanmeerbeek, Jiri Keirsse, Eva Hadadi, Jan Brughmans, Sana M. Arnouk, Ahmed E. I. Hamouda, Ayla Debraekeleer, Victor Bosteels, Yvon Elkrim, Louis Boon, Sabine Hoves, Niels Vandamme, Sofie Deschoemaeker, Sophie Janssens, Abhishek D. Garg, Martina Schmittnägel, Carola H. Ries, Damya Laoui

**Affiliations:** Myeloid Cell Immunology Lab, VIB Center for Inflammation Research, Brussels, Belgium; Lab of Cellular and Molecular Immunology, Vrije Universiteit Brussel, Brussels, Belgium; Laboratory of Cell Stress & Immunity (CSI), Department of Cellular & Molecular Medicine, KU Leuven, Leuven, Belgium; Laboratory for ER stress and Inflammation, VIB Center for Inflammation Research, Ghent, Belgium; Department of Internal Medicine and Pediatrics, Ghent University, Ghent, Belgium; JJP Biologics, Warsaw, Poland; Roche Pharmaceutical Research and Early Development, Discovery Oncology, Roche Innovation Center Munich, Penzberg, Germany; Data Mining and Modeling for Biomedicine, VIB Center for Inflammation Research, Ghent, Belgium; Department of Applied Mathematics, Computer Science and Statistics, Ghent University, Ghent, Belgium

**Keywords:** Immunotherapy, cancer, CD40 agonist, dendritic cell, macrophage, combination treatments

## Abstract

Agonistic αCD40 therapy has shown to inhibit cancer progression, but only in a fraction of patients. Hence, understanding the cancer cell-intrinsic and microenvironmental determinants of αCD40 therapy response is crucial to identify responsive patient populations and design efficient combination treatments. Here, we showed that the therapeutic efficacy of αCD40 in responder melanoma tumours, relied on pre-existing cDC1-primed CD8^+^ T cells, however cDC1s were dispensable after αCD40 administration. Surprisingly, in response to αCD40 the abundance of activated cDCs, potentially derived from cDC2s increased, thereby further activating antitumour CD8^+^ T cells. Hence, distinct cDC subsets are required to induce αCD40 responses. By contrast, lung tumours, characterised by a high abundance of macrophages, were resistant to αCD40 therapy. Combining αCD40 therapy with macrophage depletion led to tumour growth inhibition only in the presence of strong neoantigens. Accordingly, treatment with immunogenic cell-death inducing chemotherapy sensitised non-immunogenic tumours to αCD40 therapy.

## INTRODUCTION

Effective treatment of many cancer types has been consistently improving over recent decades [1]. Increased understanding of the interactions governing immune system function have led to the identification and implementation of immune checkpoint inhibitor therapies for the treatment of cancer [2,3]. Despite checkpoint inhibitors cementing themselves as invaluable therapeutic interventions, only a minority of patients experience long-term efficacy [4]. Therefore, identification of prognostic biomarkers and synergistic combination therapies that can increase the proportion of responsive patients are current focuses at the forefront of tumour immunology research [5].

Alternative therapies that aim to prime T cells rather than rescue dysfunctional T cells show great promise [6]. The TNF-receptor superfamily member CD40 is an ideal target within this context, as CD40 ligation that occurs naturally during T-cell help via CD40L results in the activation of antigen-presenting cells leading to increased T-cell priming [7–10]. Preclinical results using CD40 agonist antibodies have been shown to slow the growth of murine tumours containing strong tumour antigens [11,12], however their success in the clinic as a monotherapy was limited to a minority of melanoma patients [13]. An encouraging aspect of CD40 agonist therapy lies in the broad potential for synergistic combinations that have been shown to reduce tumour growth, including antiangiogenic therapies, tumour-associated macrophage depletion, checkpoint inhibitors, chemotherapy and radiotherapy [14–21]. While the results of these combinations are encouraging, they also hint at the importance of understanding which combination of therapies should be applied in which context.

The task of assigning synergistic combinations to an already broad landscape of different cancer (sub)types is complicated by the promiscuous expression of the CD40 receptor across multiple hematopoietic and non-hematopoietic cell types [22]. Most, but not all, antitumour effects of CD40 agonists have been shown to rely on the function of CD8^+^ T cells. However, critical cellular mediators must have activated these CD8^+^ T-cell responses. Prime candidates that have been identified as critical to CD40 efficacy are type 1 conventional dendritic cells (cDC1s) that are essential for CD8^+^ T cell priming [20,23,24]. However, studies have also implicated macrophages and other monocyte-derived cells as critical components of successful CD40 agonist-mediated antitumour immunity [25,26]. One of the most encouraging combinations investigated so far appears to involve Flt3L treatment-mediated DC boosting therapies prior to CD40 agonist therapy, with or without radiotherapy, which have been able to slow tumour growth of orthotopic and subcutaneously implanted pancreatic ductal adenocarcinoma tumours respectively [27,28].

Altogether, these results underline the importance of understanding both the cancer- and immune-specific contexture relating to successful CD40 agonist therapy. To shed further light on how the tumour microenvironments predict optimal responses to CD40 agonist therapy, and which combinatory interventions can re-sensitise non-responsive tumours, in the current study we performed single-cell RNA (scRNA-seq) sequencing on tumour-infiltrating immune cells to identify the cellular mediators of anti-CD40 (αCD40) therapy. We also utilised the *Xcr1*^wt/dtr^ mouse model to temporally deplete cDC1s and show that while the therapeutic effect of αCD40 therapy in B16F10 tumours relied on the initial function of cDC1s prior to therapy, cDC2s could be responsible for the subsequent activation, but not expansion, of antitumour T cells in response to αCD40 therapy. When comparing the αCD40-responsive B16F10 melanoma with the αCD40-resistant Lewis lung carcinoma (LLC), we identified that the highly immunosuppressive microenvironment of LLC tumours as well as their poor antigenicity limited αCD40 efficacy. By reducing suppression through αCSF1R treatment and increasing antigenicity by combination with immunogenic cell death-inducing chemotherapy, we could re-sensitise LLC tumours to αCD40 therapy.

## RESULTS

### CD40 agonist therapy repolarises B16F10 tumours resulting in reduced tumour growth

B16F10 melanoma is a frequently used preclinical mouse model in immuno-oncology that is highly infiltrated by immune cells (Fig. 1a), of which 13,0 ± 2,4% represent CD8^+^ T cells (Fig. 1b, Supplemental Fig. 1a). To assess the activation status of the tumour-infiltrating CD8^+^ T cells, we performed scRNA-seq on CD45^+^ immune cells from B16F10 tumours grown subcutaneously in C57BL/6 mice. Unsupervised clustering yielded 19 distinct clusters, which were visualised using a uniform manifold approximation and projection (UMAP) plot (Fig. 1c). The cell type of the individual immune cell clusters was identified based on their expression of canonical marker genes (Supplemental Fig. 1b). Interestingly, both defined CD8^+^ T-cell clusters in B16F10 tumours expressed high levels of genes associated with an exhausted or dysfunctional T-cell phenotype including *Pdcd1* (PD-1), *Lag3* (CD223) and *Tox* (Fig. 1d). Anti-PD-1 mAb therapy has previously been shown to reinvigorate exhausted CD8^+^ T cells, but likely due to low *Tcf7* (TCF-1) expression in the CD8^+^ T-cell population (Fig. 1d), did not result in delayed tumour growth in the B16F10 model [29,30] (Supplemental Fig. 1c), despite *Cd274* (PD-L1) gene expression within multiple different clusters (Supplemental Fig. 1d).

**Figure 1.**
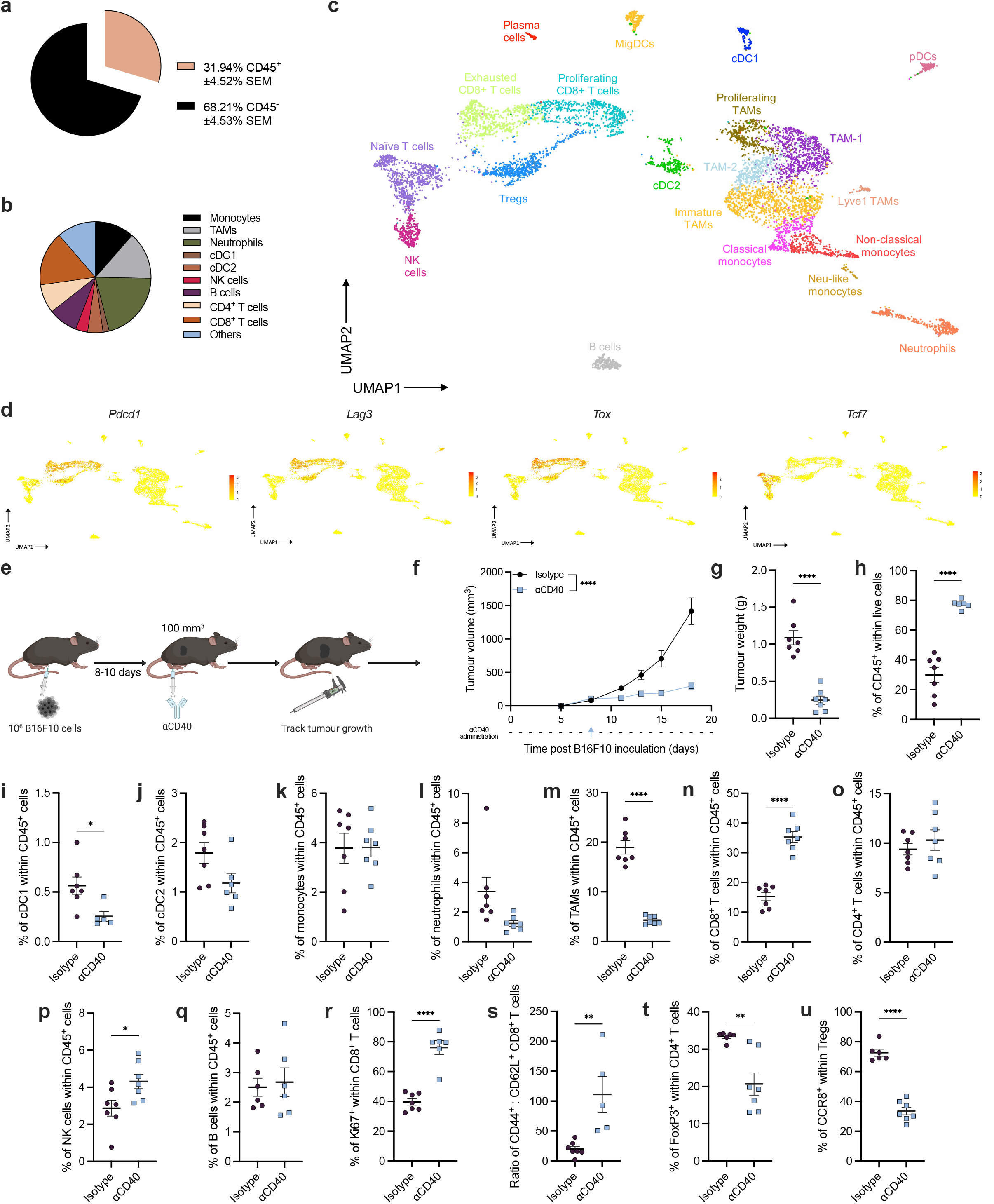
CD40 agonist therapy slows the progression of B16F10 tumours. **a**, Pie charts representing the contribution of CD45^+^ and CD45^-^ cells in B16F10 tumours and **b**, the distribution of different immune populations within the CD45^+^ fraction (averages taken from 7 individual mice). **c**, UMAP plot of 6773 CD45^+^ immune cells isolated from pools of three subcutaneous B16F10 tumours at a volume of ∼1055 ±116.4 mm^3^ and **d**, expression of several key marker genes. **e**, Schematic representation of the experimental setup indicating intraperitoneal αCD40 administration when tumours are approximately 100mm^3^ and the resulting effect of αCD40 administration on B16F10 tumour growth **(f)** and weight **(g). h**, Percentage of live cells that are CD45^+^ within isotype and αCD40 treated B16F10 tumours at day 17 post inoculation. **i-q** Frequency of multiple immune populations within isotype or αCD40 treated-day 17 B16F10 tumours. **r**, Percentage of CD8^+^ T cells that express the nuclear protein, Ki67, required for cell proliferation. **s**, Ratio of CD44^+^ CD62L^-^ effector to CD44^-^ CD62L^+^ naïve tumour infiltrating CD8^+^ T cells. **t**, Frequency of FoxP3^+^ Tregs within all tumour infiltrating CD4^+^ T cells. **u**, Percentage of Tregs within treated tumours that express CCR8. **f-v**, Representative data, shown as mean±SEM, from three independent experiments where n=7. Statistical evaluation of **f**, performed by mixed-effects analysis with Šídák’s multiple comparisons test, **g-u**, performed by unpaired t-test.

In order to investigate whether tumour growth could be arrested by targeting earlier steps in the tumour-immunity cycle, B16F10 tumour-bearing mice were treated with an anti-CD40 agonist antibody (αCD40) (Fig. 1e), as CD40 was shown to enable DC licensing and maturation resulting in subsequent priming of cytotoxic T cells [31]. CD40 agonist monotherapy significantly reduced tumour growth and weight (Fig. 1f, g) while the relative infiltration of immune cells into B16F10 tumours increased (Fig. 1h). Slightly reduced frequencies of cDC1 and cDC2 were observed 10 days after αCD40 treatment, with minor non-significant changes occurring in the monocyte and neutrophil populations (Fig. 1i, j). Tumour-associated macrophages (TAMs) were strongly decreased after successful αCD40 treatment (Fig. 1m), which is in line with previous observations showing that the presence of mature TAMs correlates with tumour size [32]. Importantly, within the lymphocytes, the abundance of cytotoxic CD8^+^ T cells was strongly increased compared to CD4^+^ T cells, NK cells and B cells (Fig. 1n-q), likely due to a higher proliferation rate (Fig. 1r). Moreover, in mice treated with αCD40, the CD8^+^ T cells displayed an effector T-cell phenotype as indicated by the increased CD44^+^ CD62L^-^ effector vs. CD44^-^ CD62L^+^ naive T-cell ratio (Fig. 1s). The elevated abundance of activated CD8^+^ T cells, was accompanied by both a decreased infiltration of FoxP3^+^ regulatory T cells (Tregs), as well as a reduced expression of CCR8 on the Tregs (Fig. 1t, u), indicative for reduced suppressive phenotype of these cells [33]. Collectively, these results indicate that CD40 agonist monotherapy is sufficient to repolarise the immune infiltrate in B16F10 tumours delaying tumour growth.

### The effect of CD40 agonist in B16F10 tumours is independent of TAMs and B cells

To investigate the mechanisms underlying the reduced tumour growth upon αCD40 agonist treatment, we first set out to determine the impact of the increased abundance and activation state of tumour-infiltrating cytotoxic CD8^+^ T cells on the inhibition of tumour progression. Systemic depletion of CD8^+^ T-cells using an αCD8 antibody restored B16F10 tumour growth in αCD40 agonist treated mice to WT levels (Fig. 2a). Next, we interrogated our scRNA-seq data to assess which cell types were expressing *Cd40* and were potentially driving anti-tumour CD8^+^ T-cell responses in B16F10 tumours. *Cd40* expression was mainly found in B cells, cDCs including cDC1s, cDC2s, CCR7^+^ DCs also termed migratory-DCs (MigDCs), DC3 [34] or mregDCs [35], and mononuclear myeloid cells including monocytes and different subsets of TAMs (Fig. 2b, Supplementary Fig. 2a). We next evaluated the expression pattern of the CD40 protein within tumour single cell suspensions via multi-colour flow cytometry. In accordance with the gene expression pattern, CD40 was only detected at the surface of Ly6C^high^ monocytes, TAMs, cDC1s, cDC2s, and B cells within B16F10 tumours (Fig. 2c, Supplementary Fig. 2b).

**Figure 2.**
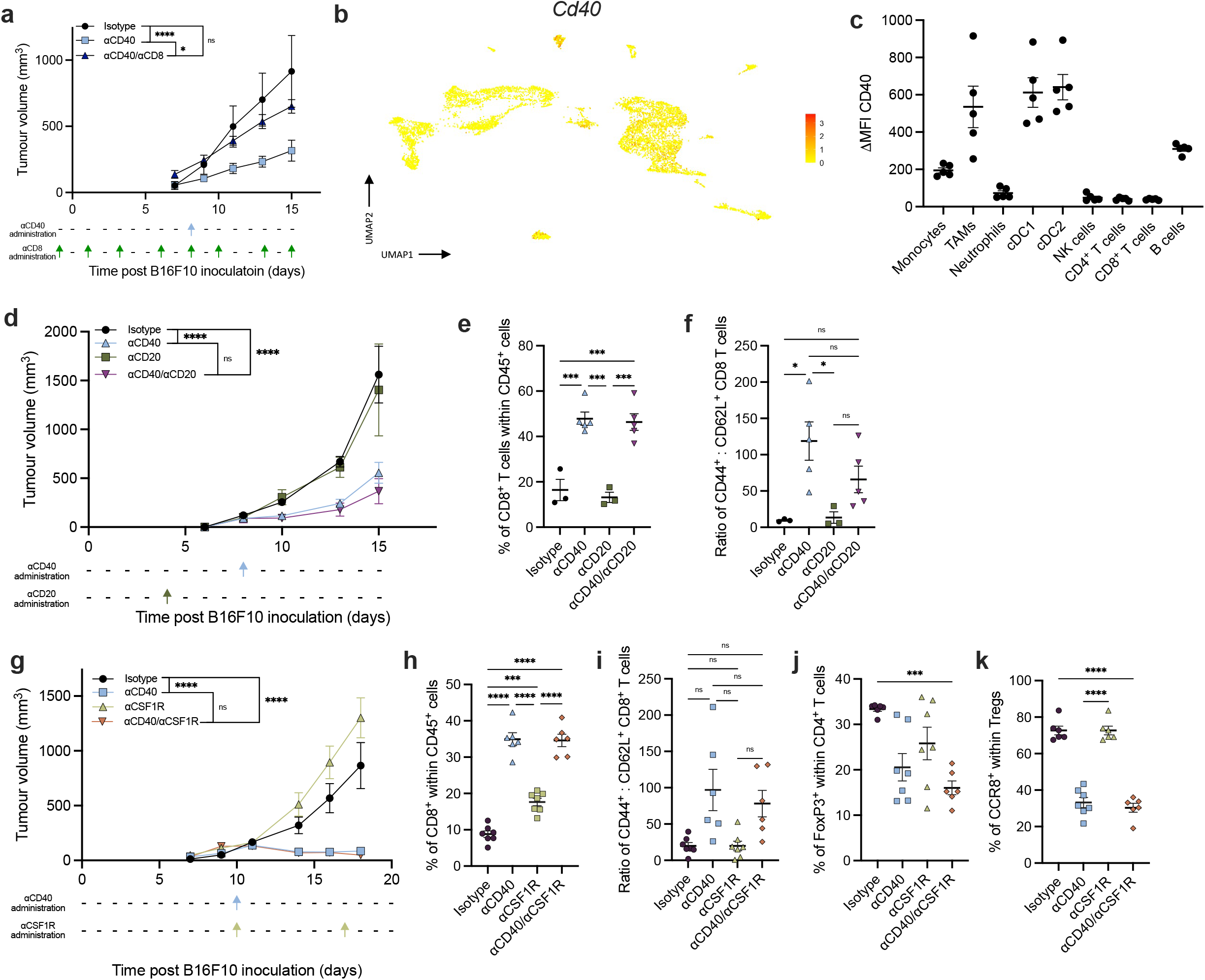
αCD40 therapy in B16F10 is TAM and B-cell independent. **a**, Growth curve of B16F10 tumours treated with combinations of isotype, αCD40 or αCD8 antibodies. (n=5). Result from one experiment **b**, UMAP showing *Cd40* mRNA expression within the CD45^+^ fraction of B16F10 tumours with volume of 1055 ±116.4 mm^3^. **c**, CD40 protein expression across distinct B16F10 tumour-infiltrating CD45^+^ cell subsets when tumours are approximately 100 mm^3^, determined by the change in median fluorescence intensity (ΔMFI) of CD40 stained samples after subtraction of FMO background signal from each sample. (n=5) Representative data from two independent experiments. **d**, Growth curve of B16F10 tumours after treatment with isotype, αCD20, αCD40 or αCD20/αCD40 antibodies (n=5) Representative of two independent experiments. **e**, Percentage of CD8^+^ T cells within day 15 B16F10 tumours after treatment. (n=5) Representative of two independent experiments. **f**, Ratio of CD44^+^ CD62L^-^ effector to CD44^-^ CD62L^+^ naïve tumour infiltrating CD8^+^ T cells. **g**, Growth curve of B16F10 tumours treated with isotype, αCD40, αCSF1R or αCD40/αCSF1R antibodies. (n=7) Representative data from three independent experiments. **h**, Percentage of CD8^+^ T cells within day 15 B16F10 tumours after treatment. **i**, Ratio of CD44^+^ effector to CD62L^+^ naïve tumour infiltrating CD8^+^ T cells. **j**, Frequency of FoxP3^+^ regulatory T cells within all tumour infiltrating CD4^+^ T cells. **k**, Percentage of Tregs within treated tumours that express CCR8. Statistical evaluation of **a**,**d**,**g**, performed by mixed-effects analysis with Tukey’s multiple comparisons test, **e**,**f**,**h**,**k** performed by ordinary one-way ANOVA with Tukey’s multiple comparisons test., **i**,**j**, performed by Brown-Forsythe and Welch ANOVA tests with Dunnett’s T3 multiple comparisons test.

To investigate whether any of these populations are involved in the therapeutic effect of αCD40 treatment, we utilised different strategies to deplete the specific immune cells including monoclonal antibodies and genetic mouse models. B cells were successfully depleted for the duration of tumour growth upon administration of 500 µg of αCD20 antibody (Supplementary Fig. 2c). The reduced tumour growth upon αCD40 therapy and increased abundance of effector CD8^+^ T cells was still unaltered in mice in which B cells were depleted (Fig. 2d-f), indicating that the reduction in tumour growth mediated by the CD40 agonist was B-cell independent. Next, to assess whether macrophages were involved in the antitumour T-cell response generated by αCD40 treatment, we depleted macrophages in tumour-bearing mice using an αCSF1R antibody (Supplementary Fig. 2d). Similarly, we could show that the anti-tumour effect of CD40 agonist therapy was independent of TAMs (Fig. 2g), as the depletion of macrophages did not revert the tumour growth, nor the increase in T-cell abundance and activation status or the decrease of immunosuppressive Tregs (Fig. 2h-k).

### The therapeutic effect of CD40 agonist in B16F10 tumours only partly relies on cDC1s

Having excluded the requirement of B cells and TAMs for the generation of a therapeutic response upon αCD40 treatment in B16F10, we next investigated the role of cDC1s. Hereto, we employed *Xcr1*^wt/dtr^ mice, which allowed temporal control of systemic cDC1 depletion upon injection of diphtheria toxin (DT) [36]. cDC1s were depleted in the tumour and in the tumour-draining lymph nodes (tdLN) 24h after DT injection (Supplementary Fig. 3a, b). Strikingly, when cDC1 depletion was initiated 24h before αCD40 administration, the therapeutic effect of the CD40 agonist therapy was unaltered (Fig. 3a), suggesting that cDC1s did not play a major role in the αCD40-mediated immune response in established B16F10 tumours. Interestingly, in *Xcr1*^wt/dtr^ mice, αCD40 treatment reduced the abundance of CD8^+^ T cells to levels comparable to isotype-treated littermate control mice (Fig. 3b), highlighting the role of cDC1 in the expansion of existing CD8^+^ T cells. Despite the inhibited expansion of CD8^+^ T cells in *Xcr1*^wt/dtr^ mice, the CD8^+^ T cells still showed an effector T-cell phenotype in *Xcr1*^wt/dtr^ mice treated with αCD40, with CD44:CD62L ratio’s similar to the T-cell phenotype in αCD40-treated littermate controls (Fig. 3c). This suggests that cDC1s are essential for CD8^+^ T-cell expansion, while other cell types are also contributed to the proper activation of existing CD8^+^ T cells into antitumour effector cells. Importantly, the depletion of these CD8^+^ T cells in *Xcr1*^wt/dtr^ mice treated with αCD40 agonist restored tumour growth in αCD40 treated mice to isotype-treated littermate control levels (Fig. 3d, Supplementary Fig. 3c). Overall, these findings suggest that therapeutic responses induced by αCD40 were driven by CD8^+^ effector T cells, independent of cDC1-mediated activation. Indeed, when cDC1s were depleted 24 hours prior to tumour inoculation and depletion was maintained throughout tumour progression, the efficacy of αCD40 agonist therapy was abrogated (Fig. 3e). Consequently, only a non-significant trend towards higher CD8^+^ T-cell levels was seen in αCD40-treated *Xcr1*^wt/dtr^ mice, which was incapable of restricting B16F10 tumour growth (Fig. 3f).

**Figure 3.**
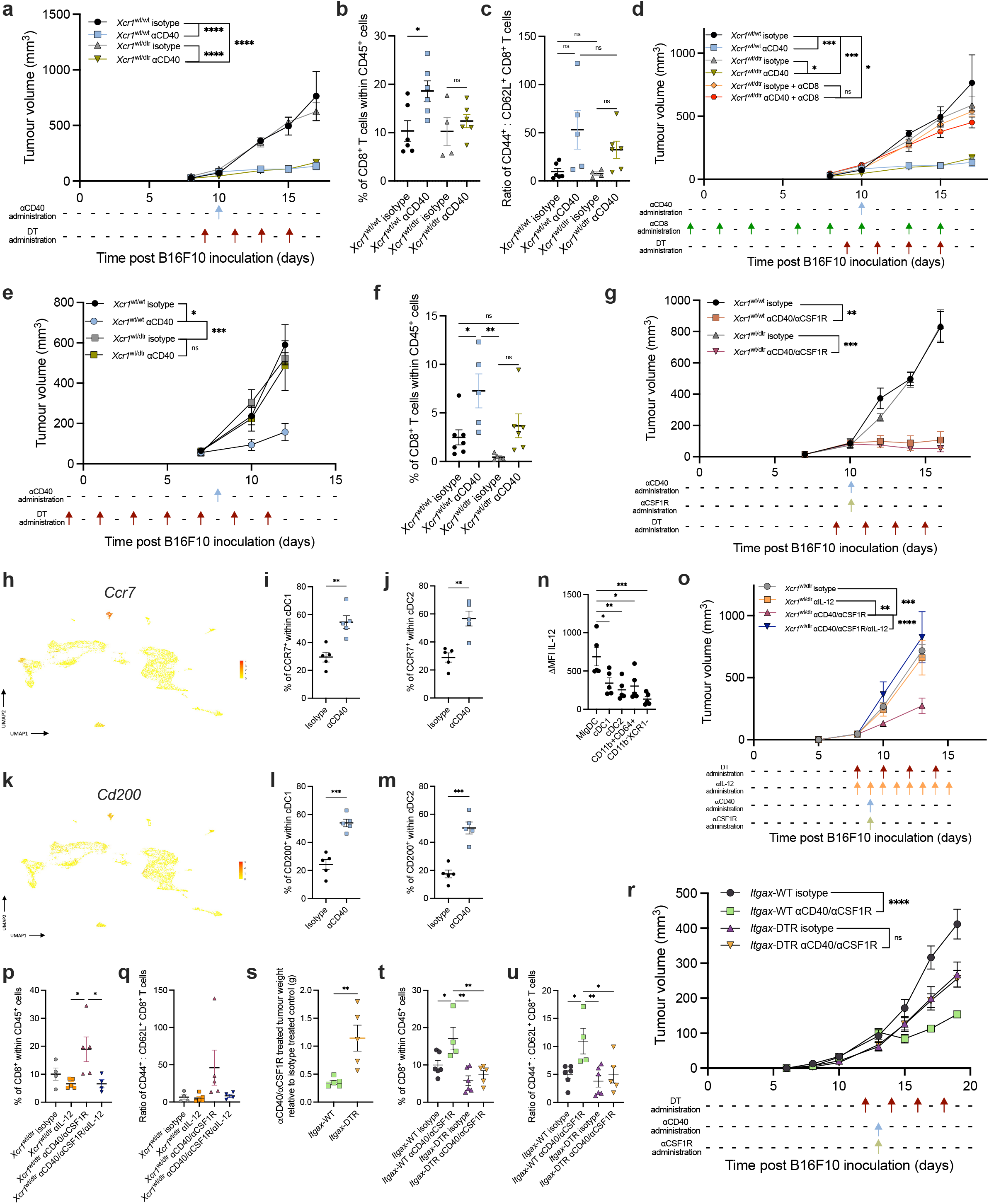
cDC1 function during early tumour growth determines αCD40 response. **a**, Growth curve of *Xcr1*^wt/wt^ and *Xcr1*^wt/dtr^ mice after treatment with isotype or αCD40 with DT administration beginning 24 hours prior to αCD40 administration, and continuing every 48 hours until the end of the experiment. (n=6) Representative data from two independent experiments. **b**, CD8^+^ T cell infiltration into day 17 B16F10 tumours from different treatment groups. **c**, Ratio of CD44^+^ effector to CD62L^+^ naïve tumour infiltrating CD8^+^ T cells. **d**, Growth curve of *Xcr1*^wt/wt^ and *Xcr1*^wt/dtr^ mice treated with isotype, αCD40, αCD8, or αCD40/αCD8 antibodies, with DT administration beginning 24 hours prior to αCD40 administration, and continuing every 48 hours until the end of the experiment. (n=6) Representative data from two independent experiments. **e**, Growth curve of *Xcr1*^wt/wt^ and *Xcr1*^wt/dtr^ mice after treatment with isotype of αCD40 with DT administration beginning 24 hours prior to B16F10 tumour implantation and continuing every 48 hours until the end of the experiment. (n=6) Representative data from two independent experiments. **f**, Infiltration of CD8^+^ T cells into treated B16F10 tumours after isotype of αCD40 administration. **g**, Growth curve of B16F10 tumours implanted in *Xcr1*^wt/wt^ and *Xcr1*^wt/dtr^ mice after isotype or αCD40/αCSF1R treatment, with DT administration beginning 24 hours prior to αCD40 administration, and continuing every 48 hours until the end of the experiment. (n=5-9), data from one experiment. **h**,**k**, UMAP plots of *Ccr7* and *Cd200* gene expression within CD45^+^ fraction of ∼1055 ±116.4 mm^3^ B16F10 tumours. **i**,**j**, Percentage of CCR7^+^ cDC1s (**i**) and cDC2s **(j)** within B16F10 tumours 24 hours after isotype or αCD40 administration. **l**,**m**, Percentage of CD200^+^ cDC1s (**l**) and cDC2s (**m**) within B16F10 tumours 24 hours after isotype or αCD40 administration. **n**, Median fluorescence intensity quantification of IL-12 expression in multiple immune subsets including MigDC, cDC1 and cDC2 after subtraction of FMO signal 24 hours after isotype or αCD40 administration. **o**, Growth curve of B16F10 tumours in *Xcr1*^wt/dtr^ mice after treatment with isotype or αCD40/αCSF1R treatment with DT administration and IL-12 neutralisation beginning 24 hours prior to αCD40/αCSF1R treatment (n=5-6), data from one experiment. **p**, CD8^+^ T-cell infiltration into day 15 B16F10 tumours after treatment with different combinations of antibodies. **q**, Ratio of CD44^+^ effector to CD62L^+^ naïve tumour infiltrating CD8^+^ T cells. **r**, Growth curve of B16F10 tumours in *Itgax-* WT and *Itgax-*DTR bone marrow chimeras after treatment with isotype or αCD40/αCSF1R with DT administration beginning 24 hours prior to αCD40/αCSF1R treatment (n=6), data from one independent experiment. **s**, Weights of tumours from *Itgax-*WT or *Itgax-*DTR mice treated with αCD40/αCSF1R relative to isotype treated mice. **t**, CD8^+^ T-cell infiltration into B16F10 tumours after treatment and based on genotype. **u**, Ratio of CD44^+^ effector to CD62L^+^ naïve tumour infiltrating CD8^+^ T cells. Statistical evaluation of **a**,**d**,**e**,**g**,**o**,**r**, performed by mixed-effects analysis and Tukey’s multiple comparisons test, **b**,**f**,**n**,**p**,**q**,**t**,**u**, performed by ordinary one-way ANOVA with Tukey’s multiple comparisons test, **c**, performed by Brown-Forsythe and Welch ANOVA tests with Dunnett’s T3 multiple comparisons test, **i**,**j**,**l**,**m**,**s**, performed by unpaired t-test. FMO, fluorescence minus one.

Next, we aimed to unravel which other antigen-presenting cells were involved in the activation of existing CD8^+^ T cells upon CD40 agonist therapy in the absence of cDC1s. Since TAMs expressed CD40 in B16F10 tumours and CD40-activated macrophages were shown to be involved in CD40-mediated tumour responses [25], we depleted TAMs in *Xcr1*^wt/dtr^ mice (Supplementary Fig. 3f). The tumour progression and activation of CD8^+^ T cells in this experiment did not differ from the results obtained in mice depleted of cDC1s 24h before αCD40 administration in which TAMs were present, suggesting that TAMs were not responsible for CD8^+^ T-cell activation in the absence of cDC1s (Fig. 3g, Supplementary Fig. 3d-f).

The only remaining immune cell types expressing CD40 that could be involved in CD8^+^ T-cell activation were cDC2s (Fig. 2b, c). Importantly, it was previously shown that the transcriptional program of cDC1s and cDC2s converged upon differentiation into MigDCs in various scRNA-seq analyses [35,37]. Two genes that showed specific upregulation in the MigDC cluster in our B16F10 data were *Ccr7* and *Cd200 (*Fig. 3h, k, Supplementary Fig. 3g). We observed that 24 hours after αCD40 administration, a higher proportion of both cDC1 and cDC2 expressed either receptor (Fig. 3i, j, l, m). When using a gating strategy that gated specifically for MigDCs based on CD200 expression (Supplementary Fig. 3h), we observed that both cDC1 and cDC2 were reduced in frequency within B16F10 tumours 24 hours after αCD40 administration while more MigDCs could be identified (Supplementary Fig. 3i-k). This is in line with the results we obtained when reanalysing a publicly available scRNA-seq dataset of murine MC38-tumour bearing WT mice generated by Zhang *et al*. [38]. Our analysis showed that the cDC1s and cDC2s were adopting a *Ccr7* expressing MigDC profile 48 hours after αCD40 treatment (Supplementary Fig. 3l, m). This might suggest that cDC2 activated by αCD40-agonist could adopt a MigDC transcriptional phenotype and mediate the activation of pre-existing CD8^+^ T-cell clones. Moreover, MHC-I levels on cDC2s were also increased upon αCD40 treatment (Supplementary Fig. 3n), further suggesting that as was shown in human [39][40], cDC2s might be able to stimulate CD8^+^ T cells.

DC-derived IL-12 was previously shown to stimulate T-cell immunity [18]. In B16F10 tumours IL-12 was mainly upregulated in the MigDC cluster, both at the transcript and protein level (Fig 3n, Supplementary Fig. 3o). To parse the role of cDC2/MigDC-derived IL-12 in effective αCD40 therapy we depleted cDC1s and TAMs and treated *Xcr1*^wt/dtr^ mice with αCD40/αCSF1R while neutralising IL-12. Blockade of IL-12 rendered the mice non-responsive to αCD40 therapy, demonstrating that IL-12 was essential to the therapeutic efficacy of αCD40 in the absence of both cDC1s and TAMs (Fig. 3o). Accordingly, the abundance and activation of tumour-infiltrating CD8^+^ T cells significantly decreased in tumours of mice that were treated with αCD40/αCSF1R/αIL-12 compared to αCD40/αCSF1R-treated mice (Fig. 3p, q). Next, to investigate whether depleting all CD11c^+^ cells (including cDC1, cDC2, and TAMs) within B16F10 tumours would abrogate the response to αCD40, we generated *Itgax-*DTR and *Itgax*-WT bone marrow chimeras to allow for continued depletion of CD11c^+^ cells. Analysis of the immune composition of these mice showed that depletion of CD11c^+^ cells was successful with strong reductions cDC1, cDC2 and TAMs within treated tumours (Supplementary Fig. 3p-r). Interestingly, we found that upon depleting CD11c^+^ cells, no differences were observed between isotype control and αCD40/αCSF1R treated mice, while tumour growth was still significantly reduced in WT reconstituted mice treated with αCD40/αCSF1R (Fig. 3r, s). Importantly, depletion of CD11c^+^ cells abrogated the increase of effector CD8^+^ T cells induced by αCD40/αCSF1R observed in control mice (Fig. 3t, u).

Overall, our data indicate that, while cDC1s play an important role in expanding CD8^+^ T cells during early phases of tumour progression, they are dispensable for the activation of existing antitumour CD8^+^ T-cell clones driving therapeutic αCD40 responses. On the other hand, our data suggests that cDC2s are capable of stimulating antitumour CD8^+^ T cells in an IL-12 dependent manner to reduce tumour growth upon CD40 agonist treatment.

### TAM depletion can further delay tumour growth after αCD40 therapy in B16F10 tumours

While αCD40 strongly reduced B16F10 tumour growth, mice were rarely tumour-free. Hence, we assessed whether αCD40 therapy could induce long lasting antitumour responses, but eventually, all mice lost tumour control approximately 5 days after the αCD40 treatment (Fig. 4a). Administration of a second dose of αCD40 five days after the first dose, did not provide any therapeutic benefit compared to mice that only received one αCD40 dose (Fig. 4a). When comparing the tumour immune infiltrate of the response phase on day 16 post tumour inoculation (tumour volume < 400 mm^3^) to the regrowth phase on day 21 post tumour inoculation (tumour volume > 600 mm^3^) upon αCD40 treatment, the myeloid compartment was more prominent in the latter at the expense of the CD8^+^ T cell-infiltrate (Fig. 4b). Moreover, there was an enrichment of MMR^+^ TAMs during the delayed regrowth phage, with MMR being a marker associated with a more protumour TAM phenotype (Fig. 4c). These data suggest that CD40 agonist therapy provides a short-term switch that polarises the TME into an immunopermissive environment, but eventually the cytotoxic response subsides, resulting in therapy resistance and tumour regrowth.

**Figure 4.**
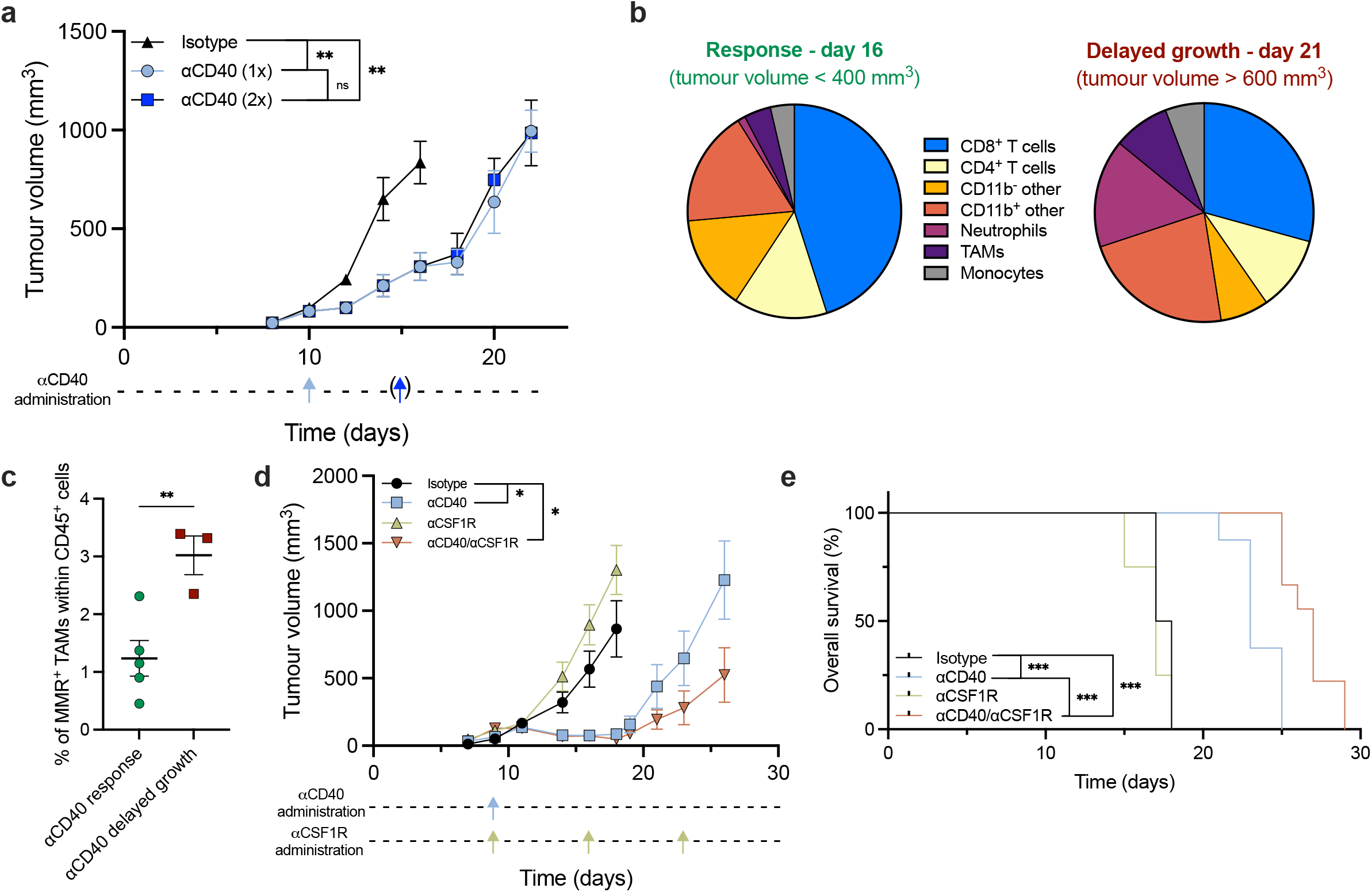
αCSF1R prolongs survival of mice after delayed B16F10 tumour regrowth. **a**, Growth curve of B16F10 implanted in WT mice after isotype, or αCD40 treatment with some mice receiving one or two doses of αCD40. (n=5-15) Representative of one experiment. **b**, Pie chart showing abundance of different immune populations within B16F10 tumours during response to αCD40 (day 16 post tumour inoculation, tumour volume of approximately 400 mm^3^), and during regrowth after αCD40 (day 21 post tumour inoculation, tumour volume of approximately 600 mm^3^) (n=5). **c**, Percentage of MMR^+^ TAMs within all CD45^+^ cells during αCD40 response and regrowth (n=3-5). **d**, Growth curve of WT B16F10 tumours until mice reach humane endpoint after treatment with isotype, αCD40, αCSF1R or αCD40/αCSF1R antibodies. (n=7) Representative data from two independent experiments. **e**, Kaplan-Meier survival curve of B16F10 tumour-bearing mice after treatment with isotype, αCD40, αCSF1R or αCD40/αCSF1R with death indicated as tumour volume surpassing > 1500mm^3^. (n=7) Representative of two independent experiments. Statistical evaluation of **a**,**d**, performed by mixed-effects analysis (using main effects only model) with Tukey’s multiple comparisons test., **c**, performed by unpaired t-test, **e**, performed by both Log-rank (Mantel-Cox) test and Gehan-Breslow-Wilcoxon test.

Given that MMR^+^ TAM have been shown to stimulate tumour relapse after therapy [41] and that in several preclinical tumour models αCD40/αCSF1R combination was able to reduce tumour growth synergistically [16,17], we wondered whether TAMs would be contributing in the delayed regrowth after αCD40 treatment. In order to test this hypothesis, we treated mice with αCD40 + αCSF1R when tumours reached 100mm^3^. Mice received one dose of αCD40, while αCSF1R was administered weekly until the humane endpoint was reached. Indeed, TAM depletion on top of αCD40 treatment significantly delayed tumour growth (Fig. 4d), resulting in a prolonged survival compared to mice which received the αCD40 as monotherapy (Fig. 4e). The TME in αCD40 + αCSF1R-treated mice contained fewer TAMs compared to αCD40-monotherapy treated tumours (Supplementary Fig. 4a, b), from which the latter included fewer TAMs that expressed Arg-1 and MMR (Supplementary Fig. 4c, d). Consequently, the abundance of CD8^+^ T cells and their effector T-cell phenotype was increased in tumours of αCD40 + αCSF1R-treated mice (Supplementary Fig. 4e, f). Depletion of CD8^+^ T cells four days after αCD40/αCSF1R administration prevented the protective effect generated by αCSF1R, showing that this effect was CD8^+^ T-cell dependent (Supplementary Fig. 4g). These results indicate that while TAM depletion was not able to further improve the therapeutic effect of αCD40 during the response phase, αCSF1R treatment could prolong the antitumour responses to CD40 agonist therapy during the delayed regrowth phase.

### B16F10 TAMs show a more immune stimulatory signature in comparison to LLC TAMs

Given the protumour role played by B16F10 TAMs upon αCD40 treatment during the delayed regrowth phase, we wondered whether the response to αCD40 would differ in preclinical models heavily infiltrated by TAMs during early tumour growth. Therefore, we utilised the Lewis lung carcinoma (LLC) model, for which we previously showed the prominence of the myeloid compartment in LLC tumours [32,42]. To investigate how the myeloid compartment differs in B16F10 compared to LLC tumours, we performed a scRNA-seq on the CD45^+^ fraction of LLC tumours at a similar tumour volumes as for the B16F10 scRNA-seq experiment. The LLC and the B16F10 tumour data were merged and reclustered jointly (Fig. 5a). After annotating the main clusters based on their expression of canonical marker genes, some major differences between the two models became obvious (Fig. 5b, Supplementary Fig. 5a). As such, the TME of LLC was characterised by a considerable heterogeneous myeloid infiltrate exemplified by expression of *Itgam*, while B16F10 tumours harboured more lymphocytes as indicated by expression of *Cd3e* (Supplementary Fig. 5b, c).

**Figure 5.**
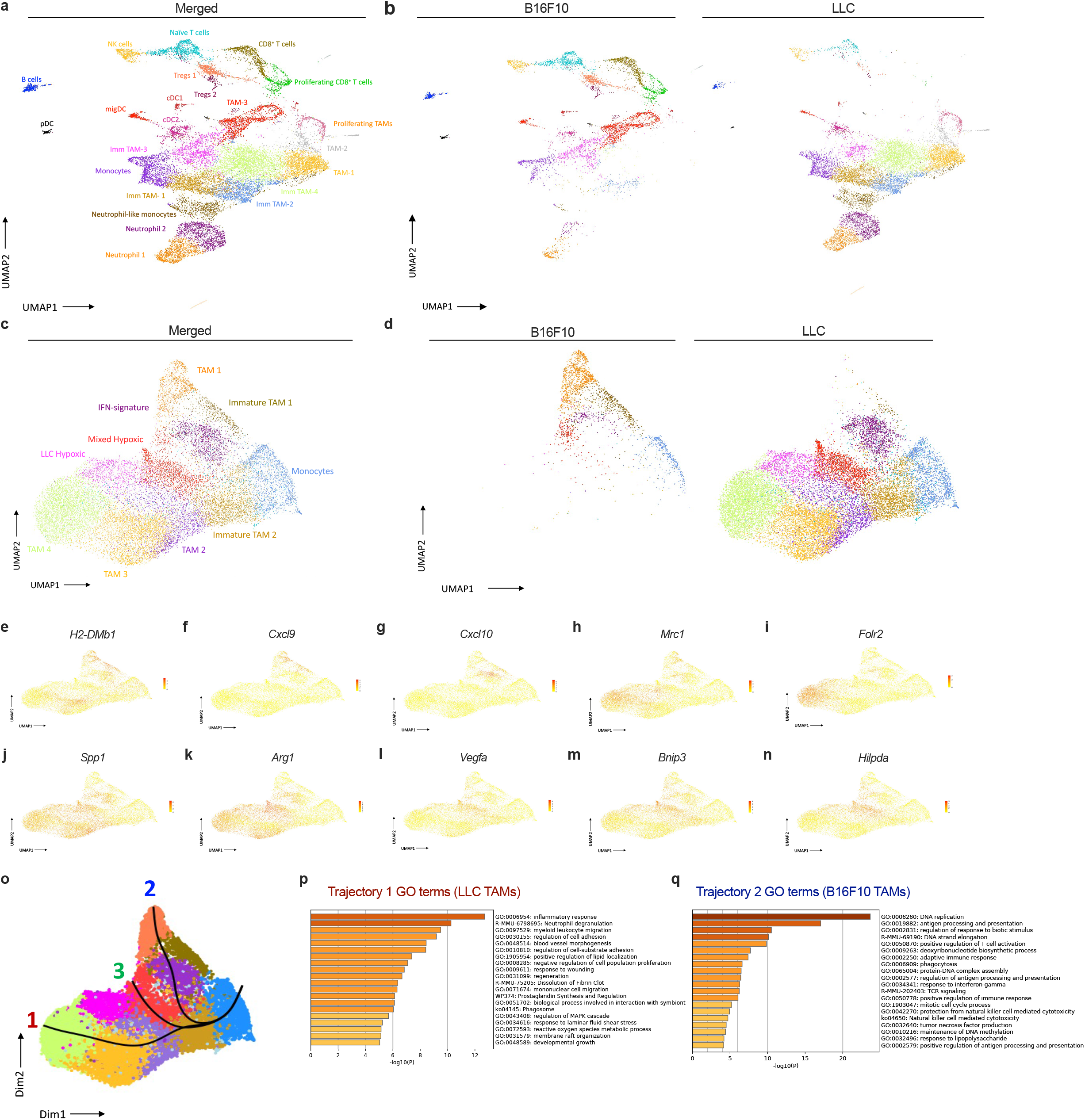
Comparison of TAM subsets in B16F10 and LLC tumours show some conserved and tumour-specific gene signatures. **a**, UMAP plot of a merged dataset, containing scRNA-seq and CITE-seq data from six individual LLC tumours (964.1 ±105.9 mm^3^) and B16F10 tumour scRNA-seq data (1055 ±116.4 mm^3^). **b**, UMAP plot of the merged dataset, comparing the individually annotated CD45^+^ cell populations between LLC and B16F10 tumours, split by tumour type. **c-d**, UMAP plots of the TAM and monocyte subset of the merged dataset, containing 18286 LLC cells and 2297 B16F10 cells, showing the identified clusters and their annotations. **e**, UMAP of the TAM and tumour-infiltrating monocyte subset of the merged dataset, containing 18286 LLC cells and 2297 B16F10 cells. **f**, UMAP of B16F10 and LLC tumour-infiltrating monocytes and TAMs separated depending on tumour of origin. **g-r**, UMAP plots showing key differentially expressed genes between the various subsets of B16F10 and LLC tumour-infiltrating monocytes and TAMs. **s**, Slingshot trajectory inference was run on the dataset containing B16F10 and LLC tumour-infiltrating monocytes and TAMs. Three distinct lineages were identified. **t-u**, Top 20 enriched gene ontology terms from a gene ontology analysis, of the genes, enriched in the endpoint of lineage 1 (LLC TAMs) and lineage 2 (B16F10 TAMs) (Wald statistic > 100, Log2FC cutoff = 1.5 and –1.5, respectively).

In order to explore the TAM heterogeneity between both models, we first subclustered the mononuclear populations, containing monocytes and TAMs and subsequently performed trajectory analysis. Some populations such as the monocyte, IFN-signature and the two hypoxic TAM clusters were represented in both tumour models, while other TAM populations such as the TAM-1 and TAM-4 clusters appeared to be unique to B16F10 or LLC tumours respectively (Fig. 5c, d). Based on differentially expressed (DE) genes between these clusters, we found that the TAMs, enriched in B16F10, expressed high levels of *H2-DMb1, Cxcl9*, and *Cxcl10* which are associated with an MHC-II^high^ M1-like inflammatory TAM phenotype. The TAM clusters enriched in LLC expressed high levels of genes associated with anti-inflammation such as *Mrc1, Folr2*, and S*pp1* (Fig 5e-j, Supplementary Fig. 5d, f). Interestingly, both LLC and B16F10 tumours harboured hypoxic TAM clusters expressing high levels of *Arg1, Vegfa, Bnip3, and Hildpa* (Fig 5k-n, Supplementary Fig. 5e).

Trajectory inference using the Slingshot method predicted 3 distinct pseudotime lineages within TAMs (Fig. 5o). Lineage 1 was mainly represented by LLC TAMs, lineage 2 by B16F10 TAMs and lineage 3, which contained the hypoxic TAMs, was shared by both models, indicating that the distinct monocyte-TAM lineages are tumour-type driven. Of note, cell percentages for each trajectory were calculated to correct for the fact that LLC tumours contained considerably more monocytes/TAMs (Supplementary Fig. 5g). Next, we performed gene ontology (GO) analysis using the Metascape analysis tool on the DE genes at the end points of lineage 1 vs lineage 2 to further unravel the divergences between TAMs from B16F10 vs LLC tumours. For the genes specific for the LLC TAM trajectory, GO analysis highlighted besides ‘inflammatory response”, mainly terms related to cell adhesion, response to wound healing, angiogenesis, and negative regulation of cell population proliferation. In contrast, the GO terms “antigen processing and presentation”, “positive regulation of T-cell activation” and “response to interferon-gamma” were highlighted for the B16F10 TAM trajectory (Fig. 5p, q). Overall, these results demonstrate that B16F10 tumours are enriched with lymphoid cells compared to LLC tumours and hint that monocyte to TAM differentiation and reprogramming is tumour-model specific with B16F10 TAMs developing toward T-cell stimulating cells, while LLC TAM develop towards potential wound-healing cells.

### LLC tumours do not respond to αCD40 therapy when combined with TAM/neutrophil depleting therapies nor therapies boosting CD8^+^ T cells

Based on the inherent differences between the B16F10 and LLC TME with the latter containing more hypoxic and tumour remodelling TAMs, we wondered whether LLC could represent a model with an inherent resistance to αCD40 therapy. Indeed, treatment of LLC tumour-bearing mice with αCD40 as a monotherapy did not reduce tumour progression (Fig. 6a, b). Given the high TAM infiltration into LLC tumours, we combined αCD40 with αCSF1R therapy, however, this resulted only in a small reduction in tumour growth (Fig. 6a-b). Nonetheless, the combination treatment slightly repolarized the remaining TAM towards an MHC-II^hi^ phenotype and increased the neutrophil, CD4^+^ and CD8^+^ T-cell infiltrate, without altering the percentages of Tregs (Fig. 6c-e, Supplementary Fig. 6a-c).

**Figure 6.**
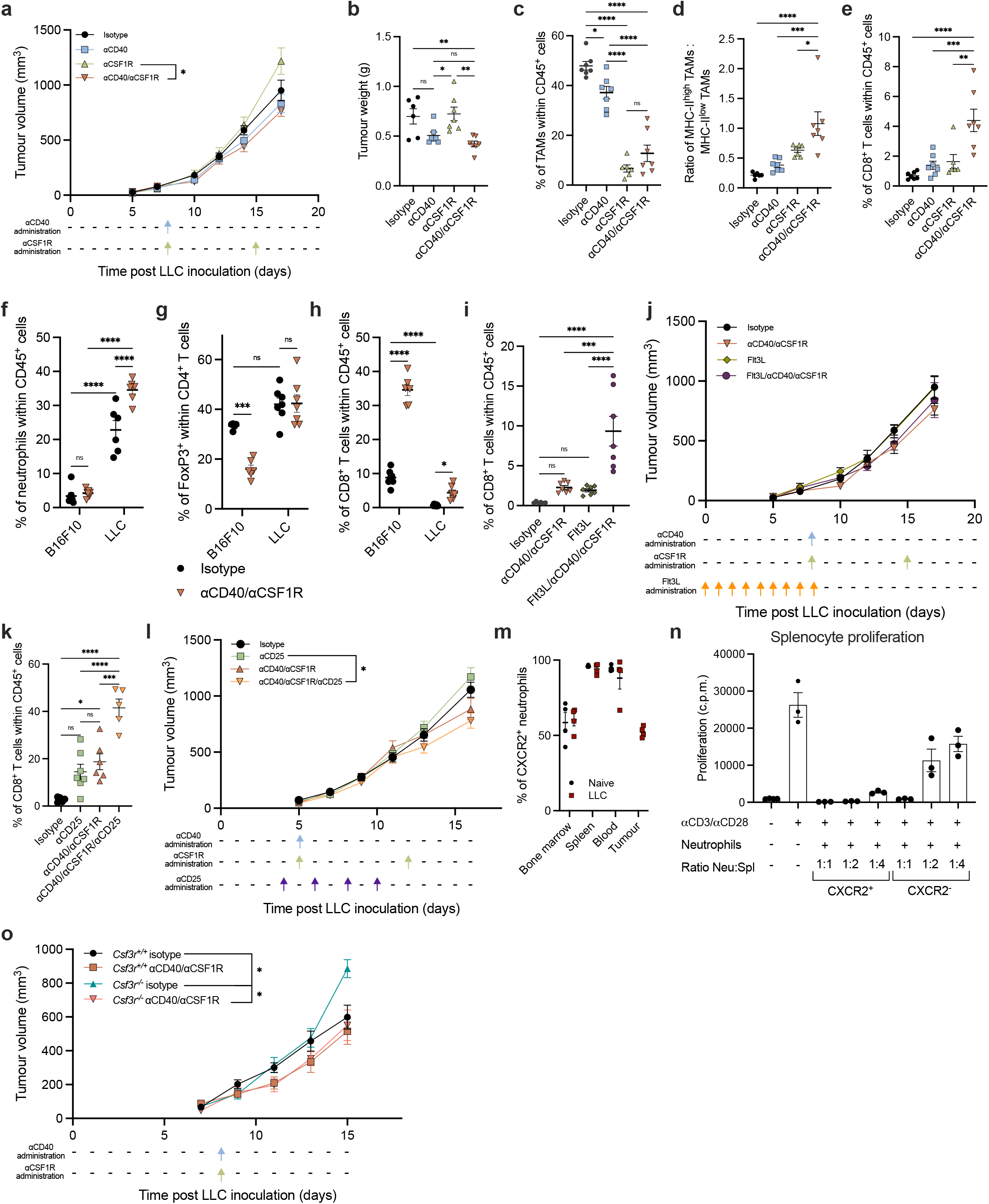
Increasing CD8^+^ T-cell infiltration into LLC tumours does not correlate with improved antitumour effects. **a**, Growth curve of LLC tumours implanted in C57Bl/6 mice after isotype, αCD40, αCSF1R, and αCD40/αCSF1R treatment. (n=7) Representative data from three independent experiments. **b**, Weights of day 17 LLC tumours from individual mice after treatment. **c**, Percentage of TAMs found within day 17 LLC tumours after treatment. **d**, Ratio of MHC-II^high^ to MHC-II^low^ TAMs within day 17 LLC tumours after isotype, αCD40, αCSF1R, or αCD40/αCSF1R treatments. **e**, Percentage of CD8^+^ T cells found within LLC tumours at day 17 post tumour implantation. **f**, Percentage of neutrophils within LLC and B16F10 tumours after isotype or αCD40/αCSF1R treatment. **g**, Percentage of FoxP3^+^ cells within CD4+ T cells of LLC and B16F10 tumours after isotype or αCD40/αCSF1R treatment. **h**, Percentage of CD8^+^ T cells within LLC and B16F10 tumours after isotype of αCD40/αCSF1R treatment. **i**, Percentage of CD8^+^ T cells within LLC tumours after pre-treatment with Flt3L and subsequent treatment with isotype or αCD40/αCSF1R antibodies. (n=7) One individual experiment. **j**, Growth curve of LLC tumours implanted in WT mice after treatment with isotype, αCD40/αCSF1R, Flt3L, or Flt3L/αCD40/αCSF1R. (n=7) one individual experiment. **k**, Percentage of CD8^+^ T cells found within LLC tumours after treatment with isotype, αCD25, αCD40/αCSF1R, or αCD40/αCSF1R/αCD25. (n=7) one individual experiment. **l**, Growth curve of LLC tumours after treatment with isotype, αCD25, αCD40/αCSF1R, or αCD40/αCSF1R/αCD25 antibodies. **m**, Percentage of neutrophils expressing the CXCR2 receptor across different tissues during naïve or at day 17 LLC tumour bearing conditions. **n**, Splenocyte proliferation following coculture of splenocytes with day 15 LLC-derived CXCR2^+^ or CXCR2^-^ neutrophils measured via 3H-thymidine incorporation (c.p.m., count per minute; n=3, data pooled from three independent experiments). **o**, Growth curve of LLC tumours implanted in *Csf3r*^+/+^ or *Csf3r*^-/-^ and treated with isotype or αCD40/αCSF1R therapy. (n=7) Representative data from two independent experiments. Statistical evaluation of **a**,**j**,**l**,**o** performed by mixed-effects analysis with Tukey’s multiple comparisons test, **b-i, k** performed by ordinary one-way ANOVA with Tukey’s multiple comparisons test.

We hypothesised that distinct immune players could be responsible for the resistance of LLC tumours towards αCD40 + αCSF1R therapy. When comparing B16F10 and LLC tumours, we observed a >4-fold increase in the abundance of tumour-infiltrating neutrophils in LLC (Fig. 6f). In addition, Tregs were strongly decreased upon αCD40 + αCSF1R treatment only in B16F10 (Fig. 6g). Both Tregs and neutrophils were shown to suppress CD8^+^ T cells in LLC tumours [33,42], which could be responsible for the lower initial abundance of CD8^+^ T cells in LLC tumours and their inability to expand upon αCD40 + αCSF1R treatment (Fig. 6h).

First, to assess whether expanding the CD8^+^ T cell number in αCD40 + αCSF1R-treated LLC would result in a therapeutic response, we employed a Flt3L treatment of LLC tumours to increase cDC numbers prior to therapy. Using an optimized Flt3L treatment schedule we were able to considerably increase intratumoural cDC numbers (Supplementary Fig. 6d, e). However, while this resulted in an increased CD8^+^ T-cell abundance in αCD40 + αCSF1R-treated mice, (Fig. 6i) tumour growth remained unaltered (Fig. 6j). Similarly, when depleting Tregs using an optimized αCD25 antibody regimen (Supplementary Fig. 6f), CD8^+^ T cells were increased upon αCD40 + αCSF1R treatment but did not result in reduced tumour growth (Fig. 6k, l).

Finally, we addressed whether neutrophils would represent a resistance mechanism to αCD40 + αCSF1R therapy. Attempts to deplete neutrophils pharmacologically in LLC tumours using αLy6G/αMAR regimens or CXCR2 inhibitors were successful, but only for a very brief window, after which neutrophils would return to normal levels in αCSF1R treatment mice (data not shown). To understand why neutrophils in LLC were not depleted using CXCR2 inhibitors, we analysed CXCR2 expression on neutrophils from bone marrow (BM), blood, spleen and tumour in naïve or LLC tumour-bearing mice. We found that in both the naïve and tumour bearing scenario, approximately 50% of the neutrophils in the BM expressed CXCR2, a receptor required for neutrophil maturation and release from the BM [43] (Fig. 6m). As expected, all neutrophils in blood and spleen expressed CXCR2, but surprisingly 50% of the neutrophils downregulated CXCR2 when reaching the TME. Interestingly, when performing a T-cell proliferation assay with CXCR2^+^ and CXCR2^-^ neutrophils, we saw that CXCR2^+^ neutrophils were more suppressive compared to CXCR2^-^ neutrophils (Fig. 6n). In αCD40 + αCSF1R-treated LLC-tumours the CXCR2^+^ neutrophil population was increased, emphasising the therapeutic potential of neutrophil depletion in αCD40 + αCSF1R-treated mice (Supplementary Fig. 6g). However, unfortunately, when using *Csf3r*^-/-^ mice, in which neutrophils are unable to egress from the BM, neutrophil depletion did not affect tumour growth of αCD40 + αCSF1R-treated mice (Fig. 6o, Supplementary Fig. 6h), implying that still other compensatory mechanisms are responsible for the therapy resistance of LLC tumours.

### Cancer cell immunogenicity determines response to αCD40/αCSF1R therapy in LLC tumours

Finally, we aimed to understand whether cancer cell intrinsic mechanisms, such as the paucity of tumour antigens, were preventing LLC-tumour bearing mice from responding to immunotherapies. Therefore, we inoculated mice with LLC cells expressing the chicken ovalbumin antigen as surrogate tumour antigen (LLC-OVA). In contrast to the results obtained in LLC, tumour growth was significantly reduced in LLC-OVA upon treatment with αCD40 and this response was even improved in combination αCD40 + αCSF1R-treated mice (Fig. 7a, b). This antitumour effect was accompanied with a strong increase in CD8^+^ T-cell abundance and a trend towards an increase in antigen specific CD8^+^ T cells, together with a repolarisation of the remaining TAMs (Fig. 7c-f). These results demonstrate that the presence of strong tumour-antigens could re-sensitise resistant models to CD40 agonist therapy.

**Figure 7.**
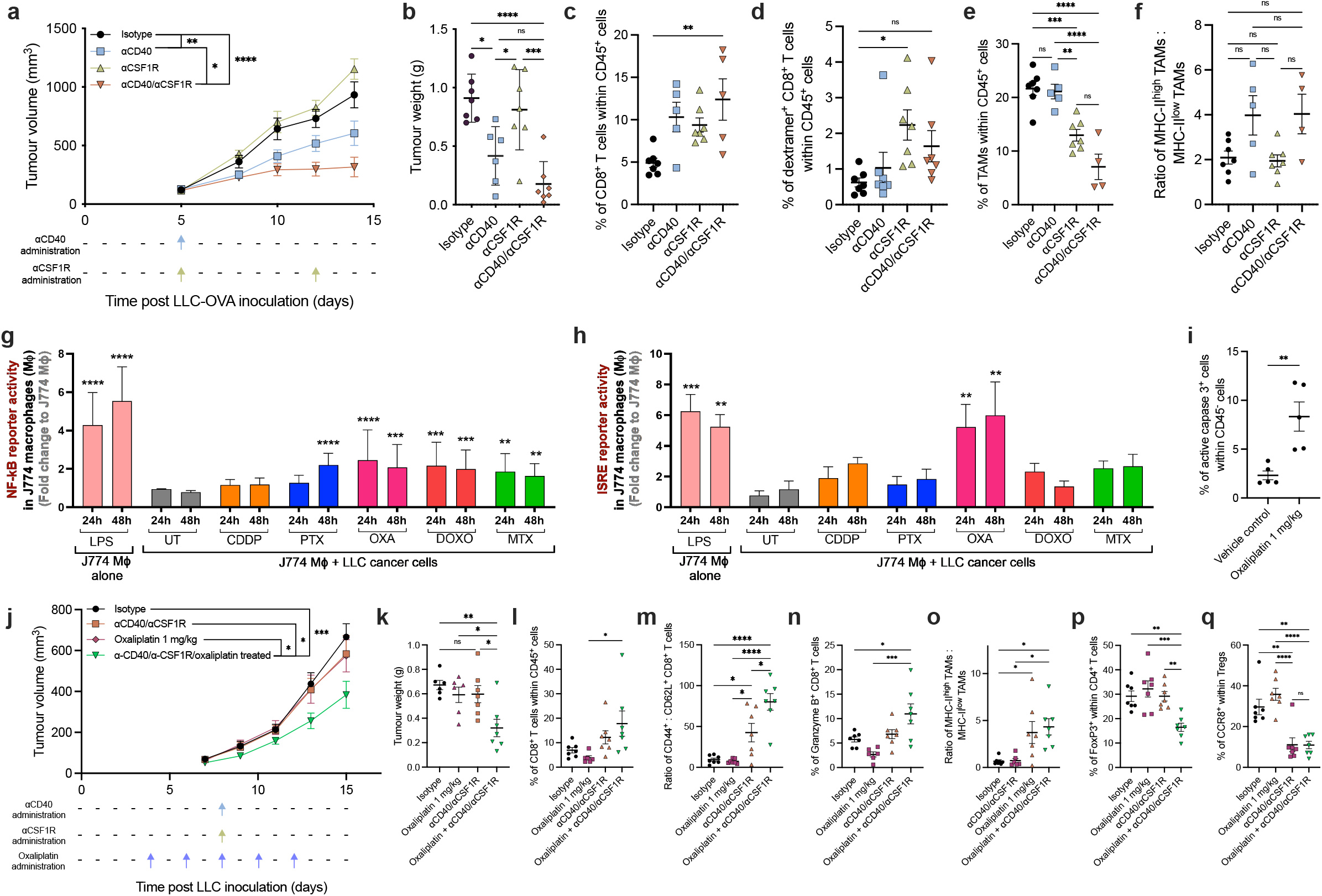
Combination with oxaliplatin synergises with αCD40/αCSF1R therapy in LLC. Growth curve of LLC-OVA tumours **(a)** and tumour weights **(b)** implanted in WT mice and treated with isotype, αCD40, αCSF1R, or αCD40/αCSF1R antibodies. (n=7) Representative data from three independent experiments. **c**, Infiltration of CD8^+^ T cells into LLC-OVA tumours after treatment with isotype, αCD40, αCSF1R, or αCD40/αCSF1R antibodies. **d**, Percentage of dextramer^+^ CD8^+^ T cells within LLC-OVA tumours after treatment with isotype, αCD40, αCSF1R, or αCD40/αCSF1R antibodies. Infiltration of TAMs into treated LLC-OVA tumours **(e)** and the ratio of MHC-II^high^ to MHC-II^low^ TAMs within treated LLC-OVA tumours **(f). g-h**, NF-kB **(g)** or ISRE **(h)** reporter activity in J774 macrophages cell 24 hours and 48 hours after culturing with LPS alone, or co-culturing with LLC cancer cells and subsequent addition of indicated chemotherapeutic compounds (n=3). **i**, Percentage of live CD45^-^ cells within day 17 LLC tumours after treatment with vehicle control or 1 mg/kg oxaliplatin. (n=5) Representative data from two independent experiments. Growth curve of LLC tumours **(j)** and corresponding tumour weights **(k)** after treatment with isotype or αCD40/αCSF1R antibodies in combination with vehicle or oxaliplatin. (n=7) Representative data from two independent experiments. **l**, Percentage of CD8^+^ T cells infiltrating day 15 LLC tumours after treatment. **m**, Ratio of CD44^+^ effector to CD62L^+^ naïve tumour infiltrating CD8^+^ T cells. **n**, Percentage of LLC tumour-infiltrating CD8^+^ T cells that express granzyme B across differently treated groups. **o**, Ratio of MHC-II^high^ to MHC-II^low^ TAMs within day 15 LLC tumours after treatment. **p**, Percentage of FoxP3^+^ cells within CD4^+^ T cells infiltrating LLC tumours after treatment. **q**, Percentage of CCR8^+^ Tregs within LLC tumours after treatment. Statistical evaluation of **a**,**j**, performed by mixed-effects analysis with Tukey’s multiple comparisons test, **b-f**,**k-q** performed by ordinary one-way ANOVA with Tukey’s multiple comparisons test, **g**,**h**, performed by two-way ANOVA, with multiple comparison correction with two-stage linear step-up procedure of Benjamini, Krieger, and Yekutieli, with a falst discovery rate at 10%, significant compared to J774+LLC UT, **i**, performed by unpaired t-test.

Next, we wanted to assess whether treating LLC tumours with immunogenic cell death (ICD)-inducing chemotherapy would recapitulate the results obtained in LLC-OVA tumours. ICD-inducers have been reported to facilitate DC-based immunogenic phagocytosis of cell corpses resulting in subsequent antigen specific T-cell activation [44]. Hereto, we first evaluated which chemotherapies could induce most potent ICD in LLC. Oxaliplatin, which is currently being used for non-small cell lung carcinoma, generated the highest NF-kB and type I interferon responses in J774 macrophages cocultured with LLC cells (Fig. 7g-h). In LLC tumour-bearing mice treated with oxaliplatin, cancer cells indeed showed increased caspase 3 activity, indicative for an increased cancer cell death (Fig. 7i). Hence, to attempt to enhance antigen uptake and presentation in LLC tumours, we combined αCD40 + αCSF1R with oxaliplatin. Oxaliplatin could significantly reduce LLC tumour progression when used in combination with αCD40 + αCSF1R (Fig. 7j-k). In addition, the proportions of CD8^+^ T cells expressing an effector phenotype and granzyme B were increased in the oxaliplatin + αCD40 + αCSF1R treated mice (Fig. 7l-n). Moreover, the remaining TAM were repolarized towards an MHC-II^high^ phenotype and less suppressive Tregs infiltrated tumours treated with the combination therapy (Fig. 7o-q).

Overall, these findings show that immunogenic cell death-inducing chemotherapy could subvert αCD40 + αCSF1R therapy resistance and thereby re-sensitise resistant tumour models.

## DISCUSSION

Cancer therapies that aim to activate a patient’s own immune cells hold a great deal of clinical promise. However, due to the potential to generate extreme adverse events, clinical application of agonist therapies must be performed with caution [45]. In the case of CD40 agonists, maximum-tolerated doses have been identified and their clinical use appears safe for patients with solid tumours [46]. Continued efforts to optimise CD40 agonist outcome will revolve around understanding optimal tumour context when αCD40 should be applied. Specific focus should attempt to identify which context-dependent cellular inputs are required for efficacy, as successful αCD40 therapy has been shown to rely on multiple cell types [16,20,25]. Finally, investigation of appropriate combinatory approaches will be beneficial to ensuring a highest possible proportion of patients can potentially benefit from αCD40 agonist therapy.

In B16F10 tumours, the involvement of CD8^+^ T cells and cDC1s was essential to generate CD40-mediated therapeutic responses. This is in accordance with previous findings which showed, using *Batf3* KO mice, that αCD40-mediated responses rely on the cDC1-CD8^+^ T-cell axis in pancreatic ductal adenocarcinoma and bladder cancer [20,47]. However, *Batf3* KO mice have two potential drawbacks. Firstly, they genetically lack critical cDC1 functions throughout all stages of tumour progression, making it challenging to properly evaluate cDC1 contribution in a temporal manner. Secondly BATF3 input has been shown recently to be critical for memory CD8^+^ T-cell development indicating that these mice may also have intrinsic issues in memory CD8^+^ T-cell formation, regardless of their dysfunctional cDC1 pool [48].

Surprisingly, using *Xcr1*^wt/dtr^ mice, we observed that cDC1 function was only essential prior to αCD40 administration in the responsive B16F10 model, and that cDC1 were redundant during the actual therapeutic phase of αCD40.

Strikingly, during the later stages of tumour progression, we showed that another cell type, likely cDC2s that upon αCD40-activation adopt a MigDC phenotype, was able to activate CD8^+^ T cells in the absence of cDC1 and TAMs. A transcriptional heterogeneity of cDC2s within mice was demonstrated in multiple cancer types, with similar counterparts also being identified in human cancers [49]. Moreover, a population of inflammatory CD64^+^ cDC2s, capable of priming both CD4^+^ and CD8^+^ T cells within a respiratory viral infection setting were recently identified [50], suggesting that cDC1s are not the only cell types able to cross present antigens to CD8^+^ T cells. As such, in the cancer context, human circulating inflammatory CD88^−^CD1c^+^CD163^+^ DCs were shown to regulate tumour immunity [51] and human cDC2s were proposed as critical mediators of cross-presentation of tumour antigens thereby promoting potent anti-tumour CTL responses [39,40]. Curiously, despite their different functional specialisations, both cDC1 and cDC2 adopt an overlapping transcriptional signature upon activation and differentiation to MigDCs [35,37,52]. Whether the ontogeny-related functions of cDC1 or cDC2 persist despite the altered signature has yet to be proven, although as cDC2 and cDC2-derived MigDCs are not depleted in *Xcr1*^wt/dtr^ mice, our data strongly suggest that these cells could be responsible for CD8^+^ T-cell activation in response to αCD40 therapy. As such, while the presence and function of intratumoural cDC1s has been shown an important player for the success of αCD40 therapy, the antitumour functions performed by cDC2s in response to CD40 agonist therapy can be more important than initially thought. However, more advanced mouse models are required to fully probe the contribution of cDC2-activated CD8^+^ T cells to αCD40 efficacy.

In response to CD40 agonist therapy, B16F10 tumour growth was controlled for multiple days. However, all mice would eventually lose tumour control and display a delayed tumour regrowth occurrence. Similar observations have been made using a combination of CD40L, TNFα and an antibody against the melanoma antigen TRP1 in which the B16F10 cancer cells formed cell-in-cell structures to avoid immune recognition [53]. This process was suggested to be mediated by IFNγ-activated CD8^+^ T cells, and once T cells are no longer present the cancer cells would disassociate from one another and continue growing. Interestingly, when we depleted TAMs once B16F10 tumours after αCD40 therapy, we observed a higher proportion of CD8^+^ T cells associated with prolonged survival and delayed tumour regrowth. This could suggest that the presence of CD8^+^ T-cell suppressive TAM could be associated with a faster disruption of the CD8^+^ T-cell-mediated cell-in-cell structures and subsequent tumour regrowth. Due to their plasticity, TAMs represent an important therapeutic target. CD40 therapy has also been shown to rely on the presence and subsequent repolarisation of TAMs to generate antitumour immunity [16,25]. The effect of αCD40 therapy on primary tumour growth was not enhanced with TAM depletion in the B16F10 model, yet TAM might have undergone a rapid reprogramming in response to CD40 agonist before their depletion as was shown in other models [16]. Interestingly, when merging scRNA-seq data of untreated B16F10 TAMs and TAMs from the heavily infiltrated LLC model, we found considerable tumour-specific heterogeneity and polarization. B16F10 TAMs appeared to be more immune-stimulatory compared to LLC TAMs. Nevertheless, when B16F10 tumours started to regrow, TAMs adopted a MMR^+^ protumour phenotype, at which point the net depletion of TAMs had a beneficial effect on survival. The broad transcriptional landscape of TAMs within tumours is being characterised thanks to modern sequencing technologies. The results of these experiments hint that a more nuanced approach to TAM depletion may result in better antitumour effects, as broad depletion using αCSF1R may also deplete antitumour TAM populations [38,42]. Although, more antitumoural TAMs have been shown to rely less on CSF1R for their survival [54].

Finally, with regard to the clinical application of CD40 agonists, our data suggest that different tumour types would benefit from different combinations of therapies. While stimulating CD8^+^ T-cell responses against tumours is a critical facet of any successful immunotherapy, our results indicate that an underlying CD8^+^ T-cell response must exist in order for αCD40 to function as a monotherapy. Therefore, if patient stratification occurs based on tumour CD3 complexity, further assessment should be made to determine whether the T cells present can recognise relevant tumour antigens or not. The intratumoural CD8^+^ T-cell pool has been shown to contain considerable irrelevant CD8^+^ T cells [55,56]. Understanding whether tumours are primarily composed of tumour reactive or bystander CD8^+^ T cells could represent a critical component for the success of αCD40 and could be informative to define appropriate complementary therapies. Patients with immune desert tumours would require more nuanced combination therapies, such as chemotherapy or radiotherapy, that would aim to generate an antitumour immune response that could be amplified with αCD40 therapy [27,28,57]. The potential risk of toxicity associated must be addressed when considering αCD40 as either a monotherapy or in combination. Encouragingly, while we suggest DCs as one of the critical mediators of antitumour immunity in response to αCD40, DCs have recently been shown to not be involved in αCD40 associated toxicity [58], suggesting that another layer of potential therapeutic combinations exist that could offset tissue damage while preserving the antitumour function of αCD40.

## MATERIALS AND METHODS

### Mouse strains

Female C57BL/6 mice were purchased from Janvier. *Xcr1*^wt/dtr^ mice were provided by Christian Kurts (University of Bonn, Germany) with the permission of Tsuneyasu Kaisho. *Csf3r*-/- mice were provided by Sebastian Jaillon and Paola Allavena (Humanitas University, Italy). CD45.1 mice were purchased from Charles River. *Itgax-*DTR mice were obtained from in-house breeding. In all experiments involving transgenic or knock-out mice, wild-type (+/+) littermate mice were used as controls as specified in the figures and figure legends.

All procedures followed the guidelines of the Belgian Council for Laboratory Animal Science and were approved by the Ethical Committee for Animal Experiments of the Vrije Universiteit Brussel (licenses 16-220-02, 18-220-19, 19-220-33, 20-220-32, 21-220-25).

### Bone marrow chimera generation

For the generation of bone marrow chimeras, female 6 week old CD45.1 mice were lethally irradiated (8 Gy). After a six hour rest period the mice were injected intravenously with 1.3×10^6^ BM cells obtained from *Itgax-*WT or *Itgax-*DTR littermate mice. The mice were used experimentally 8 weeks after BM reconstitution. Chimerism was confirmed by flow cytometry prior to tumour challenge and treatment.

### Tumour models

LLC and B16F10 cell lines (from ATCC) were cultured in DMEM (Gibco) supplemented with 10% (v/v) heat-inactivated fetal calf serum (FCS; Capricorn Scientific), 300 μg/ml L-glutamine, 100 units/ml penicillin and 100 μg/ml streptomycin. For the LLC-OVA (a kind gift from Dmitry Gabrilovich) cell line, DMEM was replaced by RPMI (Gibco).

For tumour implantation, 10^6^ LLC cells, 10^6^ B16F10 or 3×10^6^ LLC-OVA cells were injected subcutaneously into the right flank of syngeneic female C57BL/6 mice in 200 μl of HBSS.

Tumour volumes were determined by caliper measurements and calculated using the formula: V = π × (d^2^ × D)/6, where d is the shortest diameter and D is the longest diameter.

### Treatments

For CD40 agonist treatments, a single dose of 100 µg of αCD40 (clone: FGK4.5; BioXCell) agonist antibody or rat IgG2a isotype control (clone 2A3; BioXCell) was administered intraperitoneally (IP) in a volume of 100 µL HBSS when tumours reached approximately 100 mm^3^.

For macrophage depletions, 660 µg of αCSF1R (clone 2G2; provided by Roche) or murine IgG1 isotype control (clone MOPC-21; BioXCell) were administered IP in a volume of 100 µL HBSS when tumours reached approximately 100 mm^3^ with additional doses being administered weekly, if applicable.

For B-cell depletions, 500 µg of αCD20 (clone 18B12; BioXCell) or murine IgG2a (clone 2A3; BioXCell) were administered IP in a volume of 100 µL HBSS once at day 4 post tumour implantation.

To deplete cDC1s in *Xcr1*^wt/dtr^ mice, diphtheria toxin (D0564, Merck) was injected IP in *Xcr1*^wt/wt^ and *Xcr1*^wt/dtr^ mice at a dose of 25 ng/g body weight for the first dose, with following doses administered at a dose of 5 ng/g body weight.

For CD8^+^ T-cell depletions, 200 µg of αCD8 (clone YTS169; Polpharma Biologics) was administered IP in a volume of 100 µL HBSS every 2-3 days starting 1 day prior to tumour implantation.

For IL-12 neutralisation, 500 µg of αIL-12 p40 (clone C17.8, BioXCell) or rat IgG2a (clone 2A3: BioXCell) was administered IP in a volume of 100 µL HBSS daily starting 24 hours prior to αCD40 treatment and continuing until the end of the experiment.

To increase cDC numbers, 30 µg of Flt-3L-Ig (hum/hum) (clone Flt-3L Fc-G1; BioXCell) was administered IP in a volume of 50 µL HBSS every 24 hours between day 0 and day 8 post LLC tumour implantation.

To deplete Tregs, 100 µg of αCD25 (ONCC4, kindly provided by Oncurious) was administered IP in a volume of 100 µL HBSS every 48 hours between day 4 and day 10 post LLC tumour implantation, unless otherwise indicated.

Neutrophils were depleted using 75 µg αLy6G (clone 1A8; BioXCell) followed by 150 µg mouse anti-RAT (clone MAR18.5; BioXCell) administered IP in volumes of 100 µL HBSS. Alternatively, neutrophils were depleted using a CXCR2 inhibitor (SB225002, Selleck Chemicals) administered IP at a dosage of 4 mg/kg body weight.

Oxaliplatin (NSC 266046; Selleck Chemicals) was dissolved in HBSS containing 5% glucose and administered IP at 1 mg/kg body weight every 48 hours between day 4 and day 14 post LLC tumour implantation. Vehicle control (5% glucose in HBSS) was administered according to body weight of mice at time of treatment. Volumes administered were equal to 2 µL x weight of mouse (g).

### Blood collection and tissue dissociation

Blood was collected from mice in 1 ml syringes containing 0.5 mol/L EDTA. Tumours were excised, cut in small pieces, incubated with 10 U/ml collagenase I, 400 U/ml collagenase IV and 30 U/ml DNase I (Worthington) in RPMI for 20 min at 37 °C, squashed, triturated, and filtered on a 70 micron cell strainer. Spleens were mashed through a 70 micron cell strainer, bone marrow was flushed out from the femurs into RPMI. Single-cell suspensions were then treated with ACK (Ammonium-Chloride-Potassium) erythrocyte lysis buffer.

### Flow cytometry and cell sorting

Single cell suspensions were resuspended in HBSS and samples for flow cytometry analysis were incubated with Fixable Viability Dye eFluor 506 (1:1000, eBioscience) for 30 min at 4 °C. Next, cell suspensions were washed with HBSS and resuspended in HBSS with 2 mM EDTA and 1% (v/v) FCS. To prevent nonspecific antibody binding to Fcγ receptors, cells were pre-incubated with anti-CD16/CD32 (clone 2.4G2) antibody. Cell suspensions were then incubated with fluorescently labelled antibodies diluted in HBSS with 2 mM EDTA and 1% (v/v) FCS for 20 min at 4°C and then washed with the same buffer. The following fluorochrome-conjugated antibody clones were used: CD45 (30-F11), CD11b (M1/70), Ly6G (1A8), SiglecF (E50-2440), MHC-II (M5/114.15.2), Ly6C (HK1.4), F4/80 (CI:A3-1), CD11c (HL3), XCR1 (ZET), NK1.1 (PK136), CD19 (1D3), TCRβ (H57-597), CD4 (RM4-5), CD8α (53-6.7), CD44 (IM7), CD62L (MEL-14), PD-1 (RMP1-30), CCR8 (REA921), MMR (C068C2), LAG-3 (C9B7W), CXCR2 (5E8/CXCR2), CD200 (OX2), CCR7 (4B12), MHC-I (SF1-1.1), SiglecH (551), Dextramer (Immudex, Cat No. JD2163).

For intracellular staining, after extracellular staining was complete, samples were spun and fixed using the eBioscience Intracellular Fixation & Permeabilization Buffer Set (ThermoFisher, 88-8824-00) according to manufacturers instructions. The following fluorochrome-conjugated antibody clones were used: FoxP3 (FJK-16s), GZMB (GB11), Ki67 (16A8), Arg1 (14D2C43), IL-12p40 (C17.8). To measure active caspase-3 we used the FITC Active Caspase-3 Apoptosis kit (BD, 550480).

Flow cytometry data were acquired using a BD FACSCanto II (BD Biosciences) and analysed using FlowJo. The gating strategy to identify immune cell populations in tumours is shown in Supplementary Figure 1. Samples with cell contamination from the tumour-draining lymph node (identified as outliers in B cell and naive T cell abundance) were excluded from further analyses.

For fluorescence-activated cell sorting, 7AAD-CD45^+^ immune populations were sorted into RPMI containing eppendorfs for single cell sequencing. For purification of tumour-residing neutrophils, samples were enriched for CD11b^+^ cells using magnetic cell separation (Miltenyi). 7-AAD staining was used to exclude dead cells. Cell subsets were then sorted into ME medium (RPMI with 10% (v/v) FCS, 300 μg/ml L-glutamine, 100 units/ml penicillin, 100 μg/ml streptomycin, 1% (v/v) MEM non-essential amino acids (11140050, Gibco), 1 mM sodium pyruvate (Gibco) and 0.02 mM 2-mercaptoethanol (Sigma-Aldrich)). Fluorescence-activated cell sorting was performed using a BD FACSAria II (BD Biosciences).

### Single cell RNA sequencing and CITE-seq

Similarly sized tumours (collected at either day 15 or day 17 after tumour inoculation for LLC or B16F10 respectively) were pooled from three mice. The regular tissue processing procedure was followed, with the addiction of actinomycin D (Sigma-Aldrich, A1410-5MG) to each buffer. Tumour collection was performed in 30 µM, enzyme incubation and subsequent filtering in 15 µM, and all other steps in 3 µM. For scRNA-seq the single cell suspensions were stained with APC-Cy7-labelled anti-CD45 and 7AAD. Approximately 60,000 live CD45^+^ cells were sorted into ME medium using the BD FACSAria III (BD Biosciences). The sorted cells were centrifuged and resuspended in PBS (phosphate buffered saline) containing 0.04% bovine serum albumin at room temperature at an estimated final concentration of 1000 cells/µl. The CITE-seq sample was counted and 1 million cells were isolated and centrifuged. The pellet was resuspended and incubated for 30 min on ice with 25 µL of staining mix in PBS + 0.04% BSA containing APC-Cy7 labelled mouse anti-CD45 and the mouse cell surface protein antibody panel containing 174 oligo-conjugated antibodies. Subsequently the cells were washed and 60,000 live CD45^+^ cells were sorted into ME medium. Next, the 10x Genomics single-cell Bead-in Emulsions and scRNA-seq and cellular indexing of transcriptomes and epitopes by sequencing (CITE-seq) libraries were prepared as previously described [52]. The mean reads per cell for the LLC and B16F10 scRNA-seq data were 17,476 and 31,109, with a sequencing saturation metric of 38% and 42.7%, respectively. The LLC CITE-seq data yielded 11624 mean RNA reads per cell, 28.4% RNA sequencing saturation, and 2042 mean ADT reads per cell. For filtering of the low-quality cell barcodes, associated with empty droplets, the “emptyDrops” function of the DropletUtils package (v.1.8.0) has been applied on the RNA expression data, using an FDR cutoff of 0.01. The gene expression matrices were further filtered using the Scater package (v.1.16.2). The detection of outlier cells for percentage of mitochondrial genes per cell and removal of low-abundance genes were performed as previously described [59]. Library size normalization and unsupervised Leiden clustering were performed with Seurat v.3.2.3. The obtained clustering was visualized in two-dimensional scatter plots via Uniform Manifold Approximation and Projection (UMAP). Differential expression analysis was done using Wilcoxon Rank Sum test with the “FindMarkers” function of Seurat to order to identify genes, specific for each cluster. Bonferroni correction has been applied for adjustment of the p values. The processing of the ADT expression matrix was done as previously described [59].

### Trajectory inference

Trajectory inference was performed on the monocyte and TAM subsets of the mouse B16F10 and LLC tumours, using the Slingshot package (v.1.8.0) [60]. The B16F10 and the LLC datasets were merged using the “merge” command of Seurat, then monocyte and TAM clusters were subsetted and clustered using the same procedure as described above. Slingshot was run on the first 10 PCA embeddings of the monocyte/TAM subset. To identify differentially expressed genes along the identified trajectories, the package tradeseq was used (v. 1.4.0), using 5 knots for fitting the model. To find the genes that vary significantly between the two lineages, the “diffEndTest” was used, while for Identifying genes that change along a lineage, the “associationTest” function was applied [61].

### Gene ontology

To predict the putative molecular pathways and functions of the genes that distinguish the B16F10 and LLC TAMs, we performed a GO analysis on the genes that varied significantly between the two lineages using the Metascape (http://metascape.org/) online tool with default parameters [62]. We have selected the genes had Wald statistic > 100 and LogFc > 1.5 or LogFc < −1.5, respectively for the “diffEndTest” between lineage 1 and 2.

### scRNA-seq public data of αCD40 treated MC38 mice

Zhang et al. analysed CD45^+^ sorted tumours and tumour-draining lymph nodes from MC38 tumour-bearing mice treated with αCD40 antibodies [38]. We have extracted the raw FASTQ data of the day 2 treated MC38 tumours with αCD40 or isotype control (ERR3498977, ERR3499108, ERR3498975, ERR3499106, ERR3499107, ERR3499050, ERR3498978, ERR3499109, ERR3507081, ERR3507082) from https://www.ebi.ac.uk/ena/browser/view/PRJEB34105. The single cell data has been analysed as described above. DC clusters have been subsetted and reclustered.

### T-cell suppression assay

2×10^5^ neutrophils sorted from tumours were added to 2×10^5^ naïve C57BL/6 splenocytes stimulated with anti-CD3 (1 μg/ml) and anti-CD28 (2 μg/ml) and cultured in flat-bottom 96-well plates in ME medium for ex vivo cell culture described above. After 24 h of culture, 1 μCi (0.037 MBq) ^3^H-thymidine was added and after another 18 h of culture the plates were frozen and stored at −20°C after which T-cell proliferation was measured as count per minute in a liquid scintillation counter.

### NF-kB and ISRE/IRF reporter assays

The J774 macrophage-like myeloid cells were cultured in a media containing 10% heat inactivated FBS. After two passages, J774 cells containing genetic reporter constructs for detecting transcriptional sctivity of the nuclear factor kappa-light-chain-enhancer of activated B cells (NF-kB) and interferon-stimulated response element (ISRE)-binding interferon regulatory factor (IRF) were enriched via antibiotic-based selection (using 5µg/mL blasticidin and 100 µg/mL of zeocin). The J774 NF-κB and ISRE/IRF reporter myeloid cells (Invivogen) were plated with a density of 3×10^4^ cells per well in a 96-well plate. Cancer cells were plated in 10 cm dishes, and were treated with cisplatin (100 μM), paclitaxel (100 μM), doxorubicin (50 μM), mitoxantrone (0.5 μM), oxaliplatin (400 μM) or left untreated. After 24 h, the cancer cells were collected and counted. They were centrifuged at 15000 rpm for 5 min and re-suspended in J774 reporter myeloid media according to the manufacturer. These were then added on top of the J774 reporter myeloid cells, in a 1:1 ratio (in 200 μl final volume), within the 96-well plates. Stimulation with LPS (1000 ng/mL) was used as a positive control. To measure the NF-κB transcriptional activity (marked by extracellular secretion of reporter alkaline phosphatase enzyme), after 24 h or 48 h of cancer cell-J774 co-culture, 100 μl of media was transferred to a standard transparent-bottom 96-well plate. Herein, 100 μl of Quanti-BLUE substrate (Invivogen) for the alkaline phosphatase was added to each well and incubated for 4-8 h. The absorbance was measured at an optical density of 655 nm with the Biotek Synergy H1M plate reader. To measure the ISRE/IRF expression (marked by extracellular secretion of reporter luciferase enzyme), another 100μl of media was derived from the above cancer cell-J774 co-culture in a white opaque-bottom 96-well plate. Herein, 50 μl of Quanti-LUC substrate (Invivogen) for the luciferase was added and bioluminescence was directly measured with 100 millisec of signal integration, with the Biotek Synergy H1M plate reader. To account for inter-assay baseline variability a fold change to the J774 myeloid cells alone was taken from all data derived from these reporter assays.

### Schematic figures

All schematic figures were created using BioRender.com.

### Statistics

All graphs show mean±SEM. Statistical significance (p value <0.05) was determined in GraphPad Prism 9.1.2 software. For relevant pairwise comparisons, unpaired t-tests were performed. For the comparison of multiple groups, one-way analysis of variance (ANOVA) was performed, followed by a post-test. Tumour growth curves were compared by mixed-effects two-way ANOVA with multiple comparisons tests. Where appropriate statistical tests with Welch correction were performed. For statistically significant differences, the p value is indicated in graphs as the following: *p<0.05, **p<0.01, ***p<0.001, ****p<0.0001.

## DECLARATIONS

### Data and material availability

The data associated with this study are available in the main text or the supplementary materials. ScRNA-seq raw data are deposited at GEO (NCBI) under accession code xxxxx.

### Author Contribution

AM designed and performed experiments, analysed data and wrote manuscript draft. MK, HVD, JK, EH, JB, SMA, AIEH, AD, VB, YE, SD and SJ performed the experiments. DK and NVD performed or helped with bioinformatics analyses. IV and AG performed or helped with *in vitro* quantification of immunogenic cell death elicited by different chemotherapeutic compounds. LB provided materials. SH, MS, CR provided materials and funding and revised the manuscript. DL acquired funding support, supervised the study and revised the manuscript.

## Acknowledgements

We thank Ella Omasta, Marie-Therese Detobel, Nickey Van Riebeek, Nadia Abou and Christopher Stanley for technical assistance and administrative assistance. We thank Pascal. Merchiers for providing anti-CD25 antibodies. We thank Akiko Iwasaki and Orr-El Weizman for providing reagents. We would like to thank the VIB singularity platform for support and access to scRNA-seq technologies and Ria Roelandt for the library preparations. We thank Mikaël Pittet, Patrick De Baetselier, Jo Van Ginderachter and Kiavash Movahedi for insightful discussions.

## List of abbreviations

BM: bone marrow
cDC1: type 1 conventional dendritic cell
cDC2: type 2 conventional dendritic cell
DC: dendritic cell
DT: diphtheria toxin
ICD: immunogenic cell death
LLC: Lewis lung carcinoma
MigDCs: migratory DCs
scRNA-seq: single-cell RNA-sequencing
TAM: tumour-associated macrophages
tdLN: tumour-draining lymph nodes
TME: tumour microenvironment
Tregs: regulatory T cells
UMAP: Uniform Manifold Approximation and Projection

## REFERENCES

1. Jemal A, Ward EM, Johnson CJ, Cronin KA, Ma J, Ryerson AB, et al. Annual Report to the Nation on the Status of Cancer, 1975-2014, Featuring Survival. J. Natl. Cancer Inst. Oxford University Press; 2017.

2. Ishida Y, Agata Y, Shibahara K, Honjo T. Induced expression of PD-1, a novel member of the immunoglobulin gene superfamily, upon programmed cell death. EMBO J. European Molecular Biology Organization; 1992;11:3887–95.

3. Leach DR, Krummel MF, Allison JP. Enhancement of antitumor immunity by CTLA-4 blockade. Science (80-). American Association for the Advancement of Science; 1996;271:1734–6.

4. Hegde PS, Chen DS. Top 10 Challenges in Cancer Immunotherapy. Immunity. Elsevier Inc.; 2020;52:17–35.

5. Sharma P, Siddiqui BA, Anandhan S, Yadav SS, Subudhi SK, Gao J, et al. The Next Decade of Immune Checkpoint Therapy. Cancer Discov. 2021;11:838–57.

6. Vonderheide RH. CD40 Agonist Antibodies in Cancer Immunotherapy. Annu Rev Med. 2020;71:47–58.

7. Ridge JP, Di Rosa F, Matzinger P. A conditioned dendritic cell can be a temporal bridge between a CD4 + T-helper and a T-killer cell. Nature. Macmillan Magazines Ltd; 1998;393:474–8.

8. Bennett SRM, Carbone FR, Karamalis F, Flavell RA, Miller JFAP, Heath WR. Help for cytotoxic-T-cell responses is mediated by CD4O signalling. Nature. Macmillan Magazines Ltd; 1998;393:478–80.

9. Schoenberger SP, Toes REM, Van Dervoort EIH, Offringa R, Melief CJM. T-cell help for cytotoxic T lymphocytes is mediated by CD40-CD4OL interactions. Nature. Macmillan Magazines Ltd; 1998;393:480–3.

10. Van Kooten G, Banchereau J. CD40-CD40 ligand. J. Leukoc. Biol. Federation of American Societies for Experimental Biology; 2000. p. 2–17.

11. van Mierlo GJD, den Boer AT, Medema JP, van der Voort EIH, Fransen MF, Offringa R, et al. CD40 stimulation leads to effective therapy of CD40(-) tumors through induction of strong systemic cytotoxic T lymphocyte immunity. Proc Natl Acad Sci U S A [Internet]. 2002;99:5561–6. Available from: http://www.ncbi.nlm.nih.gov/pubmed/11929985

12. Mangsbo SM, Broos S, Fletcher E, Veitonmäki N, Furebring C, Dahlén E, et al. The human agonistic CD40 antibody ADC-1013 eradicates bladder tumors and generates T-cell-dependent tumor immunity. Clin Cancer Res [Internet]. 2015;21:1115–26. Available from: http://www.ncbi.nlm.nih.gov/pubmed/25316820

13. Vonderheide RH, Flaherty KT, Khalil M, Stumacher MS, Bajor DL, Hutnick NA, et al. Clinical activity and immune modulation in cancer patients treated with CP-870,893, a novel CD40 agonist monoclonal antibody. J Clin Oncol [Internet]. 2007;25:876–83. Available from: http://www.ncbi.nlm.nih.gov/pubmed/17327609

14. Kashyap AS, Schmittnaegel M, Rigamonti N, Pais-Ferreira D, Mueller P, Buchi M, et al. Optimized antiangiogenic reprogramming of the tumor microenvironment potentiates CD40 immunotherapy. Proc Natl Acad Sci U S A. 2020;117:541–51.

15. Wiehagen KR, Girgis NM, Yamada DH, Smith AA, Chan SR, Grewal IS, et al. Combination of CD40 agonism and CSF-1R blockade reconditions tumor-associated macrophages and drives potent antitumor immunity. Cancer Immunol Res. 2017;5:1109–21.

16. Hoves S, Ooi CH, Wolter C, Sade H, Bissinger S, Schmittnaegel M, et al. Rapid activation of tumor-associated macrophages boosts preexisting tumor immunity. J Exp Med. 2018;215:859–76.

17. Perry CJ, Muñoz-Rojas AR, Meeth KM, Kellman LN, Amezquita RA, Thakral D, et al. Myeloid-targeted immunotherapies act in synergy to induce inflammation and antitumor immunity. J Exp Med. 2018;215:877–93.

18. Garris CS, Arlauckas SP, Kohler RH, Trefny MP, Garren S, Piot C, et al. Successful Anti-PD-1 Cancer Immunotherapy Requires T Cell-Dendritic Cell Crosstalk Involving the Cytokines IFN-γ and IL-12. Immunity. 2018;49:1148–1161.e7.

19. Ngiow SF, Young A, Blake SJ, Hill GR, Yagita H, Teng MWL, et al. Agonistic CD40 mAb-driven IL12 reverses resistance to anti-PD1 in a T-cell-rich tumor. Cancer Res. 2016;76:6266–77.

20. Byrne KT, Vonderheide RH. CD40 Stimulation Obviates Innate Sensors and Drives T Cell Immunity in Cancer. Cell Rep [Internet]. The Author(s); 2016;15:2719–32. Available from: http://dx.doi.org/10.1016/j.celrep.2016.05.058

21. Rech AJ, Dada H, Kotzin JJ, Henao-Mejia J, Minn AJ, Victor CT-S, et al. Radiotherapy and CD40 activation separately augment immunity to checkpoint blockade in cancer. Physiol Behav. 2017;176:139–48.

22. Grewal IS, Flavell RA. CD40 and CD154 in cell-mediated immunity. Annu. Rev. Immunol. Annual Reviews 4139 El Camino Way, P.O. Box 10139, Palo Alto, CA 94303-0139, USA; 1998. p. 111–35.

23. Böttcher JP, Reis e Sousa C. The Role of Type 1 Conventional Dendritic Cells in Cancer Immunity. Trends in Cancer. Elsevier Inc.; 2018. p. 784–92.

24. Broz M, Binnewies M, Boldajipour B, Nelson A, Pollock J, Erle D, et al. Dissecting the tumor myeloid compartment reveals rare activating antigen presenting cells, critical for T cell immunity. 2015;26:B65–B65.

25. Beatty GL, Chiorean EG, Fishman MP, Saboury B, Teitelbaum UR, Sun W, et al. CD40 agonists alter tumor stroma and show efficacy against pancreatic carcinoma in mice and humans. Science (80-). Science; 2011;331:1612–6.

26. Schetters STT, Rodriguez E, Kruijssen LJW, Crommentuijn MHW, Boon L, Van Den Bossche J, et al. Monocyte-derived APCs are central to the response of PD1 checkpoint blockade and provide a therapeutic target for combination therapy. J Immunother Cancer. 2020;8:1–16.

27. Hegde S, Krisnawan VE, Herzog BH, Zuo C, Breden MA, Knolhoff BL, et al. Dendritic Cell Paucity Leads to Dysfunctional Immune Surveillance in Pancreatic Cancer. Cancer Cell. Elsevier Inc.; 2020;37:289–307.e9.

28. Lin JH, Huffman AP, Wattenberg MM, Walter DM, Carpenter EL, Feldser DM, et al. Type 1 conventional dendritic cells are systemically dysregulated early in pancreatic carcinogenesis. J Exp Med. 2020;217.

29. Kurtulus S, Madi A, Escobar G, Klapholz M, Nyman J, Christian E, et al. Checkpoint Blockade Immunotherapy Induces Dynamic Changes in PD-1 - CD8 + Tumor-Infiltrating T Cells. Immunity [Internet]. Elsevier Inc.; 2019;50:181–194.e6. Available from: https://doi.org/10.1016/j.immuni.2018.11.014

30. Siddiqui I, Schaeuble K, Chennupati V, Fuertes Marraco SA, Calderon-Copete S, Pais Ferreira D, et al. Intratumoral Tcf1+PD-1+CD8+ T Cells with Stem-like Properties Promote Tumor Control in Response to Vaccination and Checkpoint Blockade Immunotherapy. Immunity [Internet]. 2019;50:195–211.e10. Available from: http://www.ncbi.nlm.nih.gov/pubmed/30635237

31. Quezada SA, Jarvinen LZ, Lind EF, Noelle RJ. CD40/CD154 interactions at the interface of tolerance and immunity. Annu. Rev. Immunol. Annual Reviews; 2004. p. 307–28.

32. Movahedi K, Laoui D, Gysemans C, Baeten M, Stangé G, Van Bossche JD, et al. Different tumor microenvironments contain functionally distinct subsets of macrophages derived from Ly6C(high) monocytes. Cancer Res. 2010;70.

33. Van Damme H, Dombrecht B, Kiss M, Roose H, Allen E, Van Overmeire E, et al. Therapeutic depletion of CCR8+ tumor-infiltrating regulatory T cells elicits antitumor immunity and synergizes with anti-PD-1 therapy. J Immunother cancer [Internet]. 2021;9. Available from: http://www.ncbi.nlm.nih.gov/pubmed/33589525

34. Zilionis R, Engblom C, Pfirschke C, Savova V, Zemmour D, Saatcioglu HD, et al. Single-Cell Transcriptomics of Human and Mouse Lung Cancers Reveals Conserved Myeloid Populations across Individuals and Species. Immunity [Internet]. Elsevier Inc.; 2019;50:1317–1334.e10. Available from: https://doi.org/10.1016/j.immuni.2019.03.009

35. Maier B, Leader AM, Chen ST, Tung N, Chang C, LeBerichel J, et al. A conserved dendritic-cell regulatory program limits antitumour immunity. Nature. 2020;

36. Yamazaki C, Sugiyama M, Ohta T, Hemmi H, Hamada E, Sasaki I, et al. Critical roles of a dendritic cell subset expressing a chemokine receptor, XCR1. J Immunol [Internet]. 2013;190:6071–82. Available from: http://www.ncbi.nlm.nih.gov/pubmed/23670193

37. Qian J, Olbrecht S, Boeckx B, Vos H, Laoui D, Etlioglu E, et al. A pan-cancer blueprint of the heterogeneous tumor microenvironment revealed by single-cell profiling. Cell Res. Springer US; 2020;30:745–62.

38. Zhang L, Li Z, Skrzypczynska KM, Fang Q, Zhang W, O’Brien SA, et al. Single-Cell Analyses Inform Mechanisms of Myeloid-Targeted Therapies in Colon Cancer. Cell. Elsevier; 2020;181:442–459.e29.

39. Di Blasio S, Wortel IMN, van Bladel DAG, de Vries LE, Duiveman-de Boer T, Worah K, et al. Human CD1c(+) DCs are critical cellular mediators of immune responses induced by immunogenic cell death. Oncoimmunology [Internet]. 2016;5:e1192739. Available from: http://www.ncbi.nlm.nih.gov/pubmed/27622063

40. Nizzoli G, Krietsch J, Weick A, Steinfelder S, Facciotti F, Gruarin P, et al. Human CD1c+ dendritic cells secrete high levels of IL-12 and potently prime cytotoxic T-cell responses. Blood [Internet]. 2013;122:932–42. Available from: http://www.ncbi.nlm.nih.gov/pubmed/23794066

41. Hughes R, Qian BZ, Rowan C, Muthana M, Keklikoglou I, Olson OC, et al. Perivascular M2 macrophages stimulate tumor relapse after chemotherapy. Cancer Res. 2015;

42. Kiss M, Vande Walle L, Saavedra PH V, Lebegge E, Van Damme H, Murgaski A, et al. IL1β Promotes Immune Suppression in the Tumor Microenvironment Independent of the Inflammasome and Gasdermin D. Cancer Immunol Res [Internet]. 2021;9:309–23. Available from: http://www.ncbi.nlm.nih.gov/pubmed/33361087

43. Eash KJ, Greenbaum AM, Gopalan PK, Link DC. CXCR2 and CXCR4 antagonistically regulate neutrophil trafficking from murine bone marrow. J Clin Invest [Internet]. 2010;120:2423–31. Available from: http://www.ncbi.nlm.nih.gov/pubmed/20516641

44. Garg AD, Romano E, Rufo N, Agostinis P. Immunogenic versus tolerogenic phagocytosis during anticancer therapy: mechanisms and clinical translation. Cell Death Differ [Internet]. 2016;23:938–51. Available from: http://www.nature.com/articles/cdd20165

45. Suntharalingam G, Perry MR, Ward S, Brett SJ, Castello-Cortes A, Brunner MD, et al. Cytokine Storm in a Phase 1 Trial of the Anti-CD28 Monoclonal Antibody TGN1412. N Engl J Med. Massachusetts Medical Society; 2006;355:1018–28.

46. Vonderheide RH, Burg JM, Mick R, Trosko JA, Li D, Shaik MN, et al. Phase I study of the CD40 agonist antibody CP-870,893 combined with carboplatin and paclitaxel in patients with advanced solid tumors. 2013;

47. Garris CS, Wong JL, Ravetch J V, Knorr DA. Dendritic cell targeting with Fc-enhanced CD40 antibody agonists induces durable antitumor immunity in humanized mouse models of bladder cancer. Sci Transl Med [Internet]. 2021;13. Available from: http://www.ncbi.nlm.nih.gov/pubmed/34011627

48. Ataide MA, Komander K, Knöpper K, Peters AE, Wu H, Eickhoff S, et al. BATF3 programs CD8+ T cell memory. Nat Immunol. Springer US; 2020;21:1397–407.

49. Gerhard GM, Bill R, Messemaker M, Klein AM, Pittet MJ. Tumor-infiltrating dendritic cell states are conserved across solid human cancers. J Exp Med [Internet]. 2021;218. Available from: http://www.ncbi.nlm.nih.gov/pubmed/33601412

50. Bosteels C, Neyt K, Vanheerswynghels M, van Helden MJ, Sichien D, Debeuf N, et al. Inflammatory Type 2 cDCs Acquire Features of cDC1s and Macrophages to Orchestrate Immunity to Respiratory Virus Infection. Immunity [Internet]. Elsevier Inc.; 2020;52:1039–1056.e9. Available from: https://doi.org/10.1016/j.immuni.2020.04.005

51. Bourdely P, Anselmi G, Vaivode K, Ramos RN, Missolo-Koussou Y, Hidalgo S, et al. Transcriptional and Functional Analysis of CD1c+ Human Dendritic Cells Identifies a CD163+ Subset Priming CD8+CD103+ T Cells. Immunity [Internet]. 2020;53:335–352.e8. Available from: http://www.ncbi.nlm.nih.gov/pubmed/32610077

52. Pombo Antunes AR, Scheyltjens I, Lodi F, Messiaen J, Antoranz A, Duerinck J, et al. Single-cell profiling of myeloid cells in glioblastoma across species and disease stage reveals macrophage competition and specialization. Nat Neurosci [Internet]. 2021;24:595–610. Available from: http://www.ncbi.nlm.nih.gov/pubmed/33782623

53. Gutwillig A, Santana-Magal N, Farhat-Younis L, Rasoulouniriana D, Madi A, Luxenburg C, et al. Transient cell-in-cell formation underlies tumor resistance to immunotherapy. bioRxiv [Internet]. 2020;2020.09.10.287441. Available from: https://doi.org/10.1101/2020.09.10.287441

54. Van Overmeire E, Stijlemans B, Heymann F, Keirsse J, Morias Y, Elkrim Y, et al. M-CSF and GM-CSF Receptor Signaling Differentially Regulate Monocyte Maturation and Macrophage Polarization in the Tumor Microenvironment. Cancer Res [Internet]. 2016;76:35–42. Available from: http://cancerres.aacrjournals.org/cgi/doi/10.1158/0008-5472.CAN-15-0869

55. Simoni Y, Becht E, Fehlings M, Loh CY, Koo S-L, Teng KWW, et al. Bystander CD8+ T cells are abundant and phenotypically distinct in human tumour infiltrates. Nature [Internet]. 2018;557:575–9. Available from: http://www.ncbi.nlm.nih.gov/pubmed/29769722

56. Scheper W, Kelderman S, Fanchi LF, Linnemann C, Bendle G, de Rooij MAJ, et al. Low and variable tumor reactivity of the intratumoral TCR repertoire in human cancers. Nat Med [Internet]. 2019;25:89–94. Available from: http://www.ncbi.nlm.nih.gov/pubmed/30510250

57. O’Hara MH, O’Reilly EM, Varadhachary G, Wolff RA, Wainberg ZA, Ko AH, et al. CD40 agonistic monoclonal antibody APX005M (sotigalimab) and chemotherapy, with or without nivolumab, for the treatment of metastatic pancreatic adenocarcinoma: an open-label, multicentre, phase 1b study. Lancet Oncol [Internet]. 2021;22:118–31. Available from: http://www.ncbi.nlm.nih.gov/pubmed/33387490

58. Siwicki M, Gort-Freitas NA, Messemaker M, Bill R, Gungabeesoon J, Engblom C, et al. Resident Kupffer cells and neutrophils drive liver toxicity in cancer immunotherapy. Sci Immunol [Internet]. 2021;6. Available from: http://www.ncbi.nlm.nih.gov/pubmed/34215680

59. Lun ATL, McCarthy DJ, Marioni JC. A step-by-step workflow for low-level analysis of single-cell RNA-seq data with Bioconductor. F1000Research [Internet]. 2016;5:2122. Available from: http://www.ncbi.nlm.nih.gov/pubmed/27909575

60. Street K, Risso D, Fletcher RB, Das D, Ngai J, Yosef N, et al. Slingshot: Cell lineage and pseudotime inference for single-cell transcriptomics. BMC Genomics. BioMed Central Ltd.; 2018;19:477.

61. Van den Berge K, Roux de Bézieux H, Street K, Saelens W, Cannoodt R, Saeys Y, et al. Trajectory-based differential expression analysis for single-cell sequencing data. Nat Commun. Nature Research; 2020;11:1–13.

62. Zhou Y, Zhou B, Pache L, Chang M, Khodabakhshi AH, Tanaseichuk O, et al. Metascape provides a biologist-oriented resource for the analysis of systems-level datasets. Nat Commun. Nature Publishing Group; 2019;10:1–10.63. 474185v1

