## Supplementary figures for "Distinct cDC subsets co-operate in CD40 agonist response while suppressive microenvironments and lack of antigens subvert efficacy"

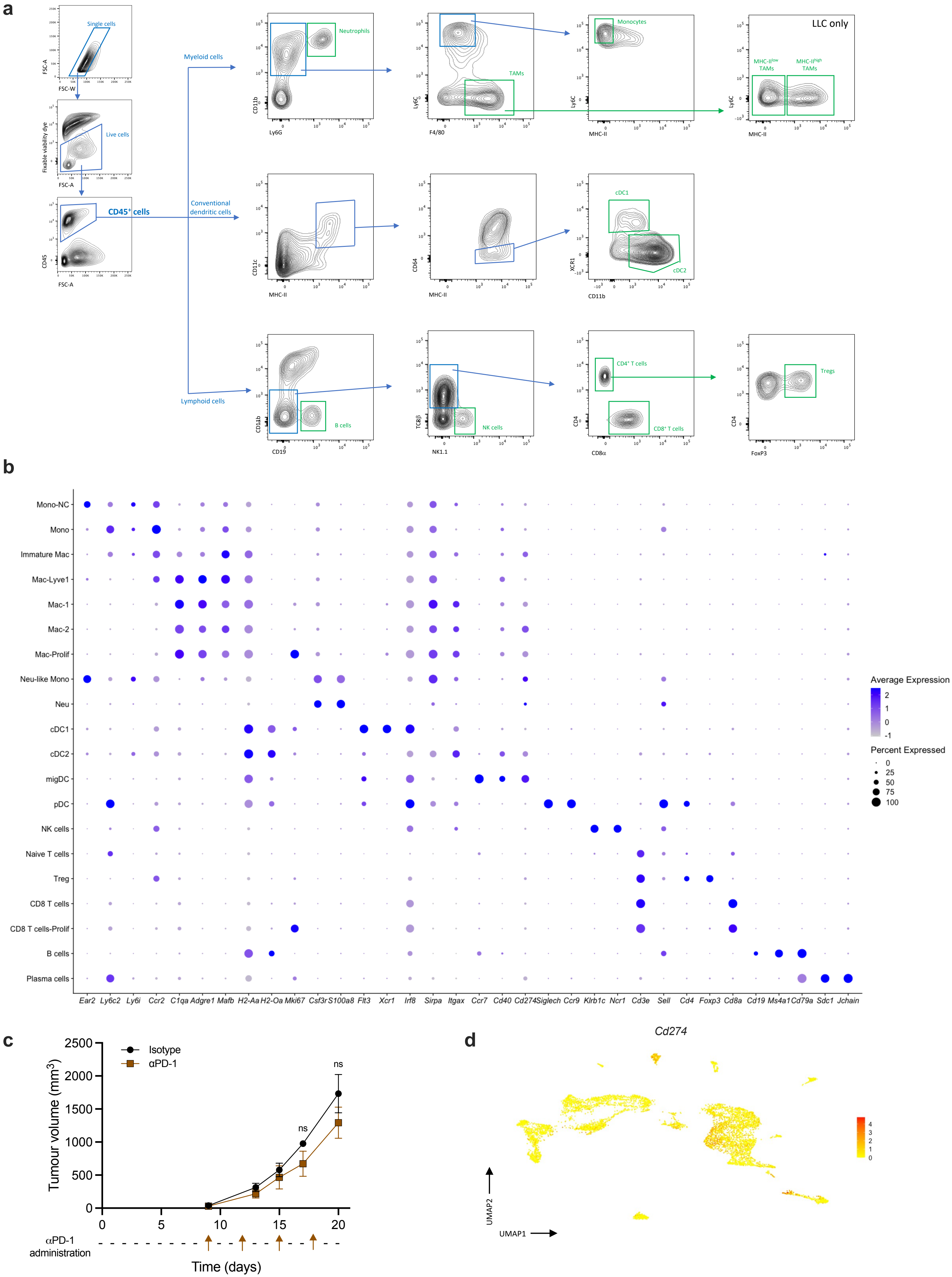

a
Single cells
FSC-A
FSC-W
Live cells
Fixable viability dye
CD45⁺ cells
CD45
Myeloid cells
Conventional dendritic cells
Lymphoid cells
Neutrophils
CD11b
Ly6G
TAMs
Ly6C
F4/80
Monocytes
MHC-II
LLC only
MHC-II low TAMs
MHC-II high TAMs
CD11c
CD64
cDC1
cDC2
XCR1
CD11b
B cells
CD19
NK cells
TCRβ
NK1.1
CD4⁺ T cells
CD8⁺ T cells
CD4
CD8α
Tregs
FoxP3
b
Mono-NC
Mono
Immature Mac
Mac-Lyve1
Mac-1
Mac-2
Mac-Prolif
Neu-like Mono
Neu
cDC1
cDC2
migDC
pDC
NK cells
Naive T cells
Treg
CD8 T cells
CD8 T cells-Prolif
B cells
Plasma cells
Ear2 Ly6c2 Ly6i Ccr2 C1qa Adgre1 Mafb H2-Aa H2-Oa Mki67 Csf3r S100a8 Flt3 Xcr1 Irf8 Sirpa Itgax Ccr7 Cd40 Cd274 Siglech Ccr9 Klrb1c Ncr1 Cd3e Sell Cd4 Foxp3 Cd8a Cd19 Ms4a1 Cd79a Sdc1 Jchain
Average Expression
2
1
0
-1
Percent Expressed
0
25
50
75
100
c
Isotype
αPD-1
Tumour volume (mm³)
2500
2000
1500
1000
500
0
ns
ns
0
5
10
15
20
αPD-1 administration
Time (days)
d
Cd274
UMAP2
UMAP1
4
3
2
1
0

**Figure S1**

**a,** Representative contour plots depicting the gating strategy used to identify individual immune cell populations within B16F10 or LLC tumours. **b,** Dot plot of selected genes that were used to identify and annotate the clusters in the B16F10 scRNA-seq dataset. The size of the dot encodes the percentage of cells within a cluster, while the colour encodes the average normalised expression level within a cluster, scaled per gene (blue is high). **c,** Growth curve of B16F10 tumour-bearing mice after IP treatment with 250 μg isotype or αPD-1 antibody. (n=3 mice per group) Representative data from two independent experiments. **d,** UMAP plot showing *Cd274* mRNA expression in the $CD45^{+}$ fraction of B16F10 tumours. Significance of **c** evaluated using mixed-effects analysis and Tukey's multiple comparisons test.

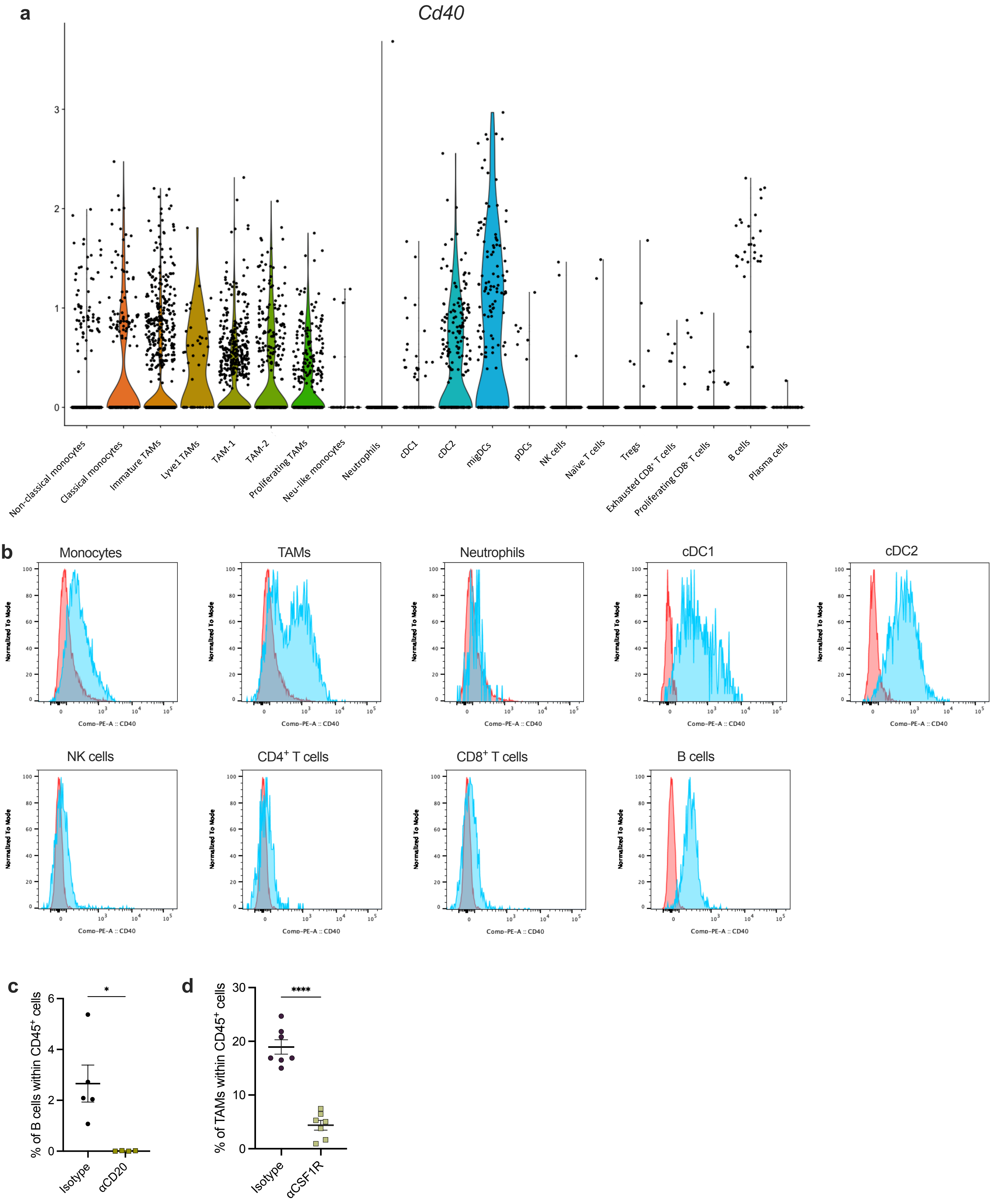
a
Cd40
0
1
2
3
Non-classical monocytes
Classical monocytes
Immature TAMs
Lyve1 TAMs
TAM-1
TAM-2
Proliferating TAMs
Neu-like monocytes
Neutrophils
cDC1
cDC2
migDCs
pDCs
NK cells
Naïve T cells
Tregs
Exhausted CD8+ T cells
Proliferating CD8+ T cells
B cells
Plasma cells
b
Monocytes
TAMs
Neutrophils
cDC1
cDC2
NK cells
CD4+ T cells
CD8+ T cells
B cells
Normalized To Mode
Comp-PE-A :: CD40
c
% of B cells within CD45+ cells
Isotype
αCD20
*
d
% of TAMs within CD45+ cells
Isotype
αCSF1R
****

**Figure S2**

**a,** Violin plot showing *Cd40* normalised mRNA expression within all $CD45^+$ immune cells clusters identified of the sequenced B16F10 tumours. **b,** Histogram plots showing CD40 expression on different immune populations via flow cytometry from 100 $mm^3$ B16F10 tumours **c,** Plot showing percentage of B cells within day 15 B16F10 tumours after isotype or aCD20 treatment on day 4. (n=5), Representative data from two independent experiments. **d,** Plot showing percentage of TAMs within day 20 B16F10 tumours after isotype or αCSF1R treatment on day 10 and 17 post tumour inoculation. (n=7) Representative data from two independent experiments. Significance of **c** and **d** was evaluated by unpaired t test *p<0.05, ****p<0.0001.

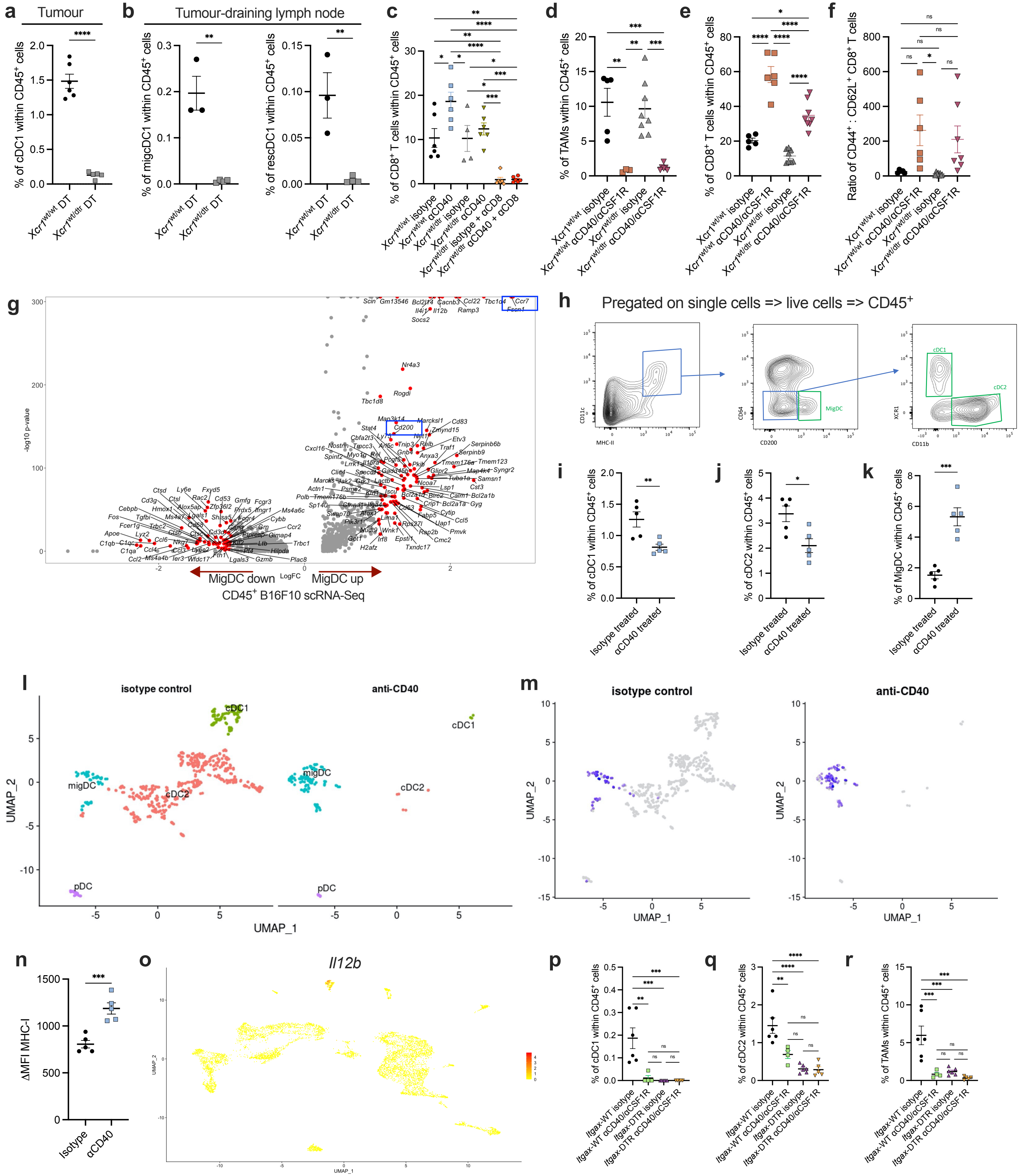

a
Tumour
% of cDC1 within CD45+ cells
Xcr1wt/wt DT
Xcr1wt/dtr DT
****
b
Tumour-draining lymph node
% of migDC1 within CD45+ cells
% of rescDC1 within CD45+ cells
**
c
% of CD8+ T cells within CD45+ cells
Xcr1wt/wt isotype
Xcr1wt/wt αCD40
Xcr1wt/dtr isotype
Xcr1wt/dtr αCD40
Xcr1wt/dtr isotype + αCD8
Xcr1wt/dtr αCD40 + αCD8
d
% of TAMs within CD45+ cells
Xcr1wt/wt isotype
Xcr1wt/wt αCD40/αCSF1R
Xcr1wt/dtr isotype
Xcr1wt/dtr αCD40/αCSF1R
e
% of CD8+ T cells within CD45+ cells
f
Ratio of CD44+ : CD62L+ CD8+ T cells
ns
g
-log10 p-value
LogFC
MigDC down
MigDC up
CD45+ B16F10 scRNA-Seq
Nr4a3
Rogdi
Tbc1d8
Cd200
Ccr7
Fscn1
Il12b
Socs2
h
Pregated on single cells => live cells => CD45+
CD11c
MHC-II
CD64
CD200
MigDC
XCR1
CD11b
cDC1
cDC2
i
% of cDC1 within CD45+ cells
Isotype treated
αCD40 treated
j
% of cDC2 within CD45+ cells
k
% of MigDC within CD45+ cells
l
isotype control
anti-CD40
cDC1
migDC
cDC2
pDC
UMAP_1
UMAP_2
m
isotype control
anti-CD40
UMAP_1
UMAP_2
n
ΔMFI MHC-I
Isotype
αCD40
o
Il12b
UMAP_1
UMAP_2
p
% of cDC1 within CD45+ cells
Itgax-WT isotype
Itgax-WT αCD40/αCSF1R
Itgax-DTR isotype
Itgax-DTR αCD40/αCSF1R
q
% of cDC2 within CD45+ cells
r
% of TAMs within CD45+ cells

**Figure S3**

**a,** Percentage of cDC1 within B16F10 tumours in *Xcr1*$^{wt/wt}$ and *Xcr1*$^{wt/dtr}$ mice 24 hours after DT administration (n=6) Representative data from two independent experiments. **b,** Percentage of migratory (left) and resident (right) cDC1 in tumour-draining lymph nodes of *Xcr1*$^{wt/wt}$ and *Xcr1*$^{wt/dtr}$ mice 24 hours after DT administration. (n=3) Representative data from two independent experiments. **c,** Percentage of CD8$^{+}$ T cells within B16F10 tumours isolated from *Xcr1*$^{wt/wt}$ and *Xcr1*$^{wt/dtr}$ mice after treatment with DT in combination with isotype, αCD40, αCD8 or αCD40/αCD8 antibodies. (n=6) Representative data from two independent experiments. **d,** Percentage of TAMs within B16F10 tumours isolated from *Xcr1*$^{wt/wt}$ and *Xcr1*$^{wt/dtr}$ after treatment with DT in combination with isotype or αCD40/αCSF1R treatment. (n=5-9) Data from one experiment. **e,** Percentage of CD8$^{+}$ T cells within B16F10 tumours isolated from *Xcr1*$^{wt/wt}$ and *Xcr1*$^{wt/dtr}$ after treatment with DT in combination with isotype or αCD40/αCSF1R treatment. (n=5-9) Data from one experiment. **f,** Ratio of CD44$^{+}$ CD62L$^{-}$ effector to CD44$^{-}$ CD62L$^{+}$ naïve tumour infiltrating CD8$^{+}$ T cells within B16F10 tumours isolated from *Xcr1*$^{wt/wt}$ and *Xcr1*$^{wt/dtr}$ after treatment with DT in combination with isotype or αCD40/αCSF1R treatment. **g,** Volcano plot showing genes that are up- and down- regulated in the migDC population in comparison to all other clusters within CD45$^{+}$ fraction B16F10 tumours. **h,** Gating strategy used for examining cDC1 and cDC2 with specific gating for MigDCs using CD200. **i,j,k,** Abundance of MigDC, cDC1, and cDC2 within 100 mm$^{3}$ B16F10 tumours 24 hours after isotype or αCD40 administration, using gatings based on CD200 expression by MigDCs. (n=5) Representative data from two independent experiments. **l,** UMAP plot of the DC populations from a public scRNA-seq dataset of CD45$^{+}$ sorted MC38 tumours 48 hours after isotype or αCD40 antibody administration. The UMAP plot is separated by treatment. **m,** UMAP plot of the DC populations from CD45$^{+}$ sorted MC38 tumours 48 hours after isotype or αCD40 antibody administration, showing *Ccr7* gene expression and separated by treatment. **n,** Expression of MHC-I on B16F10 tumour-residing cDC2s 24 hours after isotype or aCD40 treatment. **o,** UMAP plot showing *Il12b* mRNA expression in the $CD45^+$ fraction of B16F10 tumours. **p-r,** Abundance of cDC1 (**p**), cDC2 (**q**), and TAMs (**r**) in B16F10 tumours from *Itgax*-WT and *Itgax*-DTR bone marrow chimeras treated with DT and subsequent treatment with aCD40/aCSF1R or isotype control antibodies. (n=6), data from one experiment. Significance of **a, b, i-k, n** was evaluated by unpaired t test. Significance of **c-f, p-r** was evaluated by ordinary one-way ANOVA with Tukey's multiple comparisons test. *p<0.05, **p<0.01, ***p<0.001, ****p<0.0001.

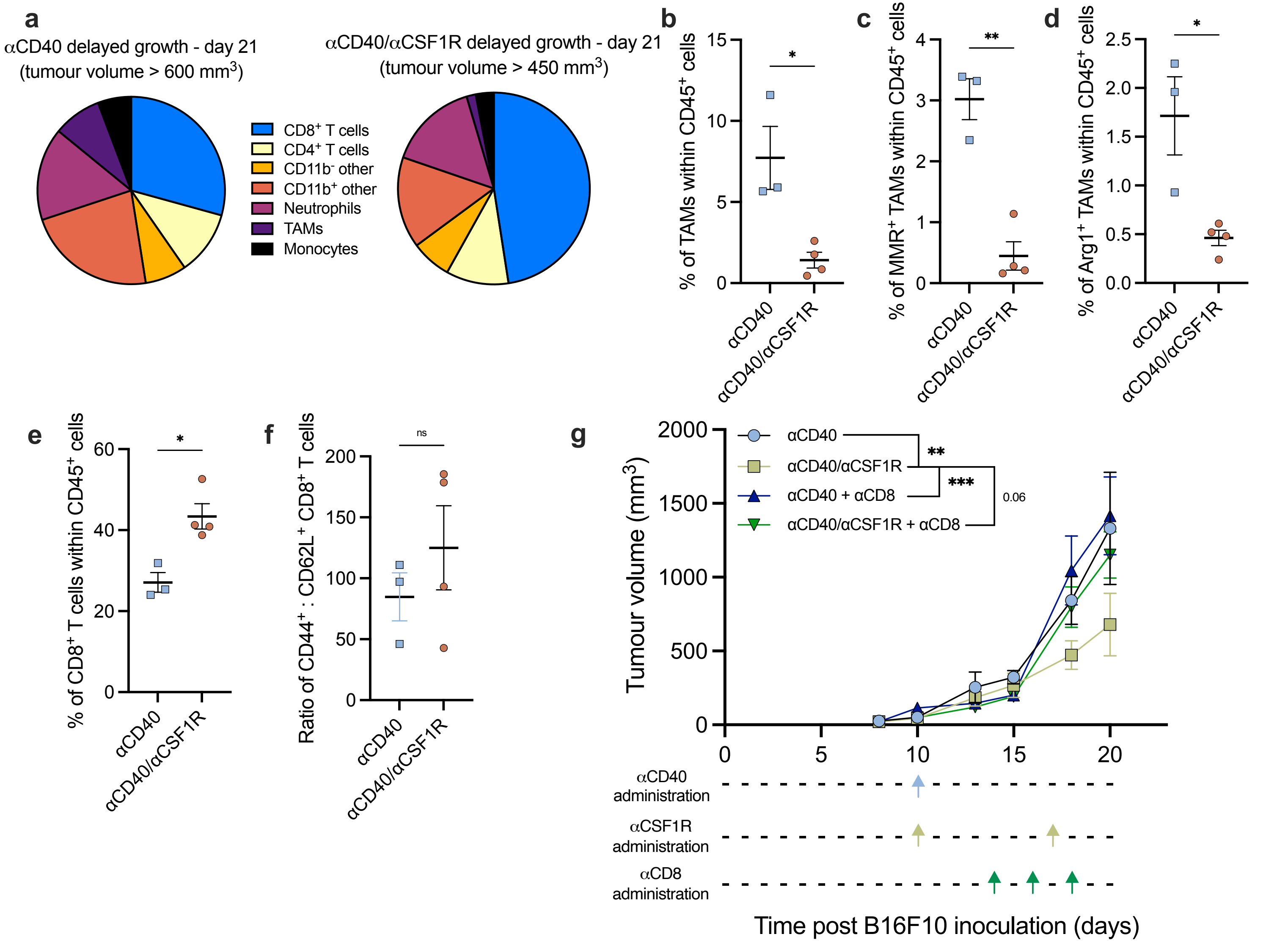
a
αCD40 delayed growth - day 21
(tumour volume > 600 mm3)
αCD40/αCSF1R delayed growth - day 21
(tumour volume > 450 mm3)
CD8+ T cells
CD4+ T cells
CD11b- other
CD11b+ other
Neutrophils
TAMs
Monocytes
b
% of TAMs within CD45+ cells
c
% of MMR+ TAMs within CD45+ cells
d
% of Arg1+ TAMs within CD45+ cells
e
% of CD8+ T cells within CD45+ cells
f
Ratio of CD44+ : CD62L+ CD8+ T cells
ns
αCD40
αCD40/αCSF1R
g
Tumour volume (mm3)
αCD40
αCD40/αCSF1R
αCD40 + αCD8
αCD40/αCSF1R + αCD8
**
***
0.06
αCD40 administration
αCSF1R administration
αCD8 administration
Time post B16F10 inoculation (days)

**Figure S4**

**a,** Pie charts indicating the contribution of different immune populations to the CD45$^+$ fraction of B16F10 tumours after delayed regrowth upon αCD40 or αCD40/αCSF1R. **b,** Percentage of TAMs within WT B16F10 tumours after treatment of αCD40 or αCD40/αCSF1R. **c,** Percentage of MMR$^+$ TAMs within CD45$^+$ cells of B16F10 tumours after αCD40 or αCD40/αCSF1R treatment. **d,** Percentage of Arginase1$^+$ TAMs within CD45$^+$ cells from B16F10 tumours after αCD40 or αCD40/αCSF1R treatment. **e,** Percentage of CD8$^+$ T cells within B16F10 tumours after αCD40 or αCD40/αCSF1R treatment. **f,** Ratio of CD44$^+$ effector to CD62L$^+$ naïve tumour infiltrating CD8$^+$ T cells after αCD40 or αCD40/αCSF1R treatment. **g,** Growth curve of B16F10 in WT mice treated with αCD40 or αCD40/αCSF1R in which αCD8$^+$ T cells were depleted as from 4 days after αCD40 administration. (n=6) Data from one experiment. **b-f,** (n=3-5) Representative data from two independent experiments. Significance was evaluated by unpaired t test. Significance of **g** was calculated using 2way ANOVA with Tukey's multiple comparisons test. *$p<0.05$, **$p<0.01$, ***$p<0.001$.

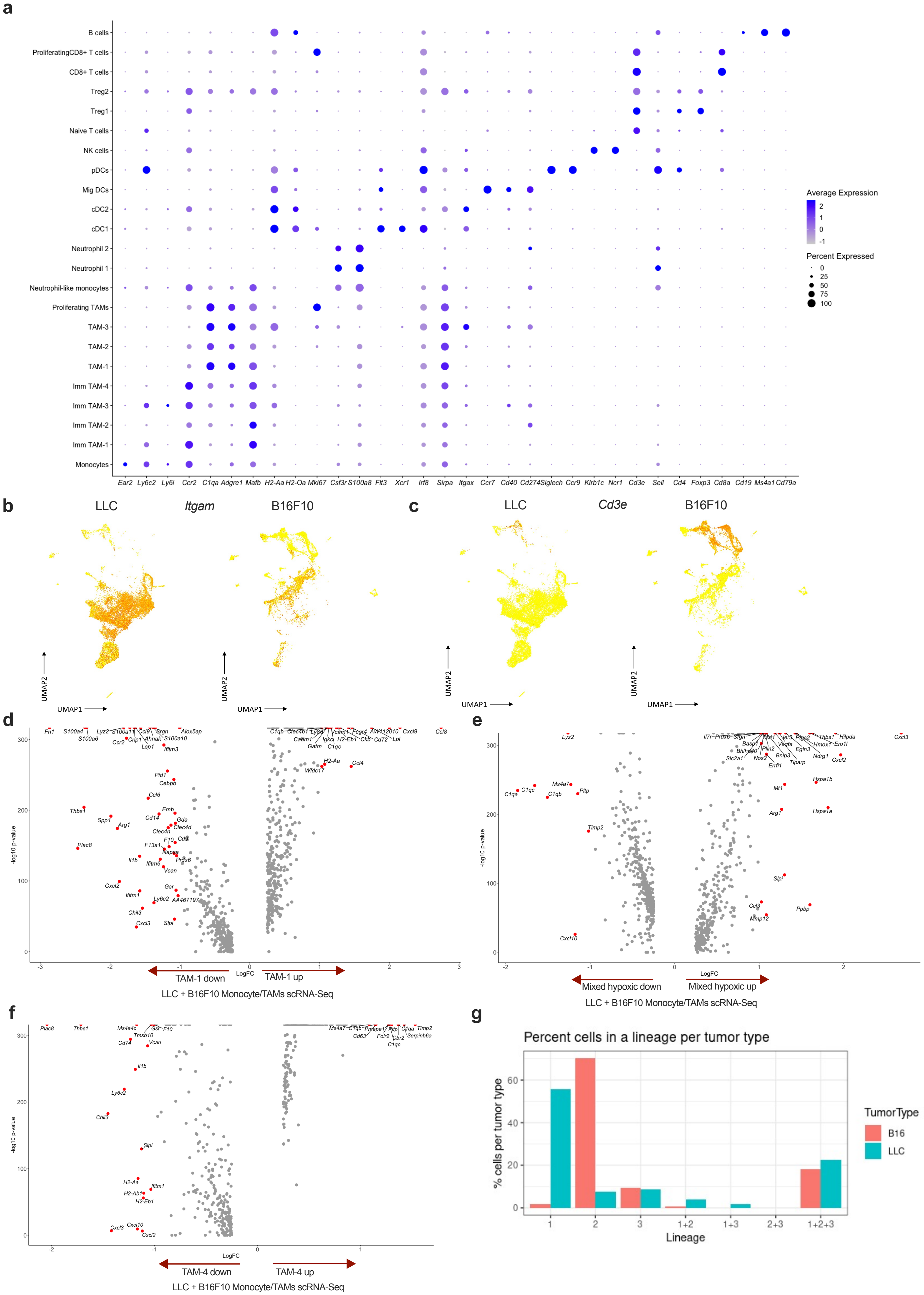
a
B cells
ProliferatingCD8+ T cells
CD8+ T cells
Treg2
Treg1
Naive T cells
NK cells
pDCs
Mig DCs
cDC2
cDC1
Neutrophil 2
Neutrophil 1
Neutrophil-like monocytes
Proliferating TAMs
TAM-3
TAM-2
TAM-1
Imm TAM-4
Imm TAM-3
Imm TAM-2
Imm TAM-1
Monocytes
Ear2 Ly6c2 Ly6i Ccr2 C1qa Adgre1 Mafb H2-Aa H2-Oa Mki67 Csf3r S100a8 Flt3 Xcr1 Irf8 Sirpa Itgax Ccr7 Cd40 Cd274 Siglech Ccr9 Klrb1c Ncr1 Cd3e Sell Cd4 Foxp3 Cd8a Cd19 Ms4a1 Cd79a
Average Expression
2
1
0
-1
Percent Expressed
0
25
50
75
100
b
LLC
Itgam
B16F10
UMAP2
UMAP1
c
LLC
Cd3e
B16F10
UMAP2
UMAP1
d
-log10 p-value
LogFC
TAM-1 down
TAM-1 up
LLC + B16F10 Monocyte/TAMs scRNA-Seq
e
-log10 p-value
LogFC
Mixed hypoxic down
Mixed hypoxic up
LLC + B16F10 Monocyte/TAMs scRNA-Seq
f
-log10 p-value
LogFC
TAM-4 down
TAM-4 up
LLC + B16F10 Monocyte/TAMs scRNA-Seq
g
Percent cells in a lineage per tumor type
% cells per tumor type
Lineage
1
2
3
1+2
1+3
2+3
1+2+3
TumorType
B16
LLC

**Figure S5**

**a,** Dot plot of selected genes that were used to identify and annotate the clusters in the merged B16F10 and LLC dataset. UMAP plot**,** showing *Itgam* (**b)** and *Cd3e* (**c**) expression, separated by tumour type. **d,** Volcano plot showing the DE genes of TAM-1 cluster in comparison to all other clusters in the monocyte/TAM subset of the B16F10/LLC dataset. **c,** Volcano plot showing the DE genes of mixed hypoxic cluster in comparison to all other clusters in the monocyte/TAM subset of the B16F10/LLC dataset. **d,** Volcano plot showing the DE genes of TAM-4 cluster in comparison to all other clusters in the monocyte/TAM subset of the B16F10/LLC dataset. **e,** Percentage of cells from each sample that comprise each of the distinct lineages, detected by Slingshot trajectory inference analysis in the monocyte/TAM subset of the B16F10/LLC dataset.

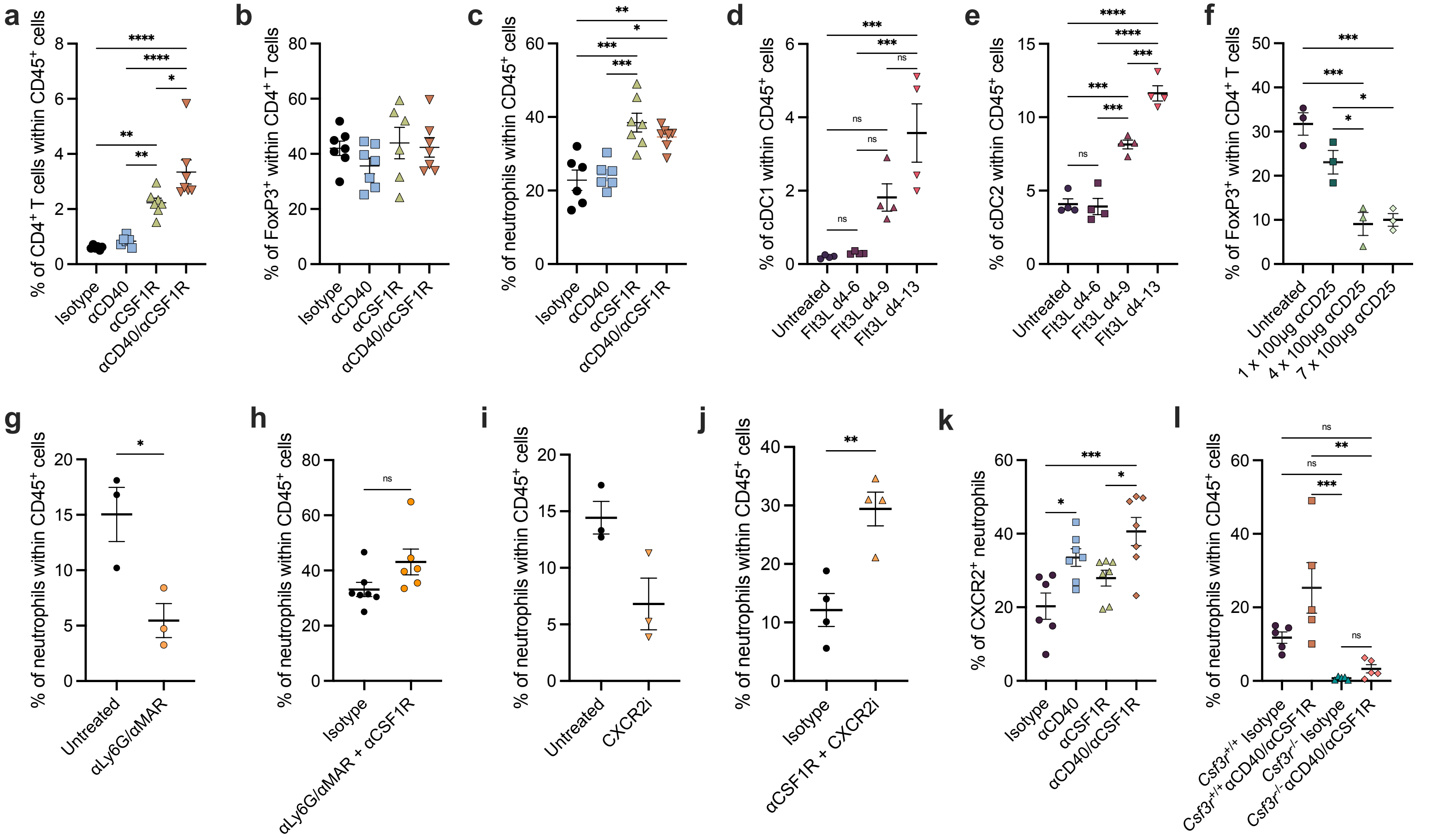
a
% of CD4+ T cells within CD45+ cells
Isotype
αCD40
αCSF1R
αCD40/αCSF1R
b
% of FoxP3+ within CD4+ T cells
c
% of neutrophils within CD45+ cells
d
% of cDC1 within CD45+ cells
Untreated
Flt3L d4-6
Flt3L d4-9
Flt3L d4-13
e
% of cDC2 within CD45+ cells
f
% of FoxP3+ within CD4+ T cells
1 x 100μg αCD25
4 x 100μg αCD25
7 x 100μg αCD25
g
αLy6G/αMAR
h
αLy6G/αMAR + αCSF1R
i
CXCR2i
j
αCSF1R + CXCR2i
k
% of CXCR2+ neutrophils
l
Csf3r+/+ Isotype
Csf3r+/+ αCD40/αCSF1R
Csf3r-/- Isotype
Csf3r-/- αCD40/αCSF1R

**Figure S6**

**a,** $CD4^+$ T cell infiltration into LLC tumours after treatment with isotype, αCD40, αCSF1R, or αCD40/αCSF1R antibodies. **b,** Percentage of $CD4^+$ T cells that express the FoxP3 transcription factor across treatment groups. **c,** Neutrophil infiltration into LLC tumours after treatment with isotype, αCD40, αCSF1R, or αCD40/αCSF1R antibodies. **a-c,** (n=7) Representative data from three independent experiments. Percentage of cDC1s **(d)** and cDC2s **(e)** within LLC tumours after treatment with 30 µg of Flt3L for different durations during tumour growth. (n=4) Representative data from three independent experiments. **f,** Percentage of $FoxP3^+$ cells within $CD4^+$ T cells identified within LLC tumours after different treatment regimens using aCD25 targeting antibodies. (n=3) Data from one experiment. **g,** Percentage of neutrophils within day 15 LLC tumours after treatment with isotype or αLy6G/αMAR antibodies on d11, d12, d13 and d14 post tumour inoculation. (n=3) Representative data from two independent experiments. **h,** Percentage of neutrophils within day 17 LLC tumours after IP administration with αCSF1R on d10 and IP administration of αLy6G/αMAR on d11, d12, d13, d14, d15, d16 after tumour inoculation. (n=7) Data from one experiment. **i,** Percentage of neutrophils within d17 LLC tumours after IP treatment with vehicle control or CXCR2i beginning day 4 post tumour inoculation. (n=3) Data from one experiment. **j,** Percentage of neutrophils within d17 LLC tumours after treatment with isotype or αCSF1R together with CXCR2i. (n=4) Data from one experiment. **k,** Percentage of neutrophils expressing CXCR2 within LLC tumours after treatment with isotype, αCD40, αCSF1R, or αCD40/αCSF1R antibodies. (n=7) Data from one experiment. **l,** Percentage of neutrophil contribution to $CD45^+$ tumour infiltrating cells in $Csf3r^{+/+}$ or $Csf3r^{-/-}$ and treated with isotype or αCD40/αCSF1R therapy. (n=7), Representative data from two independent experiments. Significance of **a-f** and **k, l** was evaluated by ordinary one-way ANOVA with Tukeys' multiple comparisons test. Significance of **g-j** was evaluated by unpaired t test. *p<0.05, **p<0.01, ***p<0.001, ****p<0.0001 by ordinary one-way ANOVA with Tukey's multiple comparisons test.
